# Human Cytomegalovirus Infection Reduces an Endogenous Antiviral Fatty Acid by Promoting Host Metabolism

**DOI:** 10.1101/2025.03.31.646481

**Authors:** Yuecheng Xi, Rebekah L. Mokry, Nicholas D. Armas, Ian Kline, Maxwell Wegner, John G. Purdy

## Abstract

Some viruses, including human cytomegalovirus (HCMV), induce the synthesis of fatty acids and lipids to ensure that the lipid environment of infected cells supports virus replication. HCMV infection broadly reprograms metabolism to ensure central carbon metabolism provides the metabolites required for anabolic synthesis of nucleotides, proteins, and lipids while also meeting the energy demands placed on the infected cells. While HCMV infection increases the levels of most very long chain fatty acids (VLCFA), we found that the levels of erucic acid (EA), a C22:1 monounsaturated VLCFA, are reduced. Treating infected cells with EA disrupted a late step in virus replication, resulting in the release of virions with reduced infectivity. Moreover, we used lipidomics to determine that EA-treated cells had elevated levels of lipids containing a combination of a C22:1 tail and a VLC polyunsaturated fatty acid tail (VLC-PUFA). We demonstrate that fatty acid elongase 5 (ELOVL5) mediated production of VLC-PUFAs is stimulated by HCMV infection. ELOVL5 aided the increase in lipids with C22:1 plus VLC-PUFA tails following EA treatment and reduced the overall level of C22:1 in HCMV-infected cells. Moreover, we found that ELOVL5 mollified EA inhibition of HCMV replication, suggesting ELOVL5 plays a critical role in reducing the level of an endogenous FA with antiviral properties. Our study provides insight into how infection may increase the synthesis of an antiviral metabolite or FA and how the virus may evade their antiviral effect by promoting their metabolism.

## Introduction

Lipids and fatty acids (FAs) are essential building blocks for viral infection, including human cytomegalovirus (HCMV). HCMV is a herpesvirus that infects 60–90% of the population worldwide [1, 2]. While life-long persistent infections are typically asymptomatic in healthy individuals, the virus causes life-threatening illnesses in immunocompromised people, including organ transplant recipients and cancer patients [1, 2]. HCMV is also a leading cause of congenital infection and developmental disabilities [3–5]. Current treatments include inhibitors of the viral DNA polymerase, such as ganciclovir [6]. However, the current treatments are poor options due to low bioavailability or toxicity [7]. Resistance mutations to some drugs occur in several HCMV genes, further limiting the current treatment options [6].

Since viruses do not encode a complete metabolic network, they rely on host cells to provide for their metabolic needs during replication. The dependency on host metabolism is a weakness that the host can exploit to suppress viruses like HCMV. Following infection, HCMV induces lipid synthesis, elevating the levels of those with very long chain fatty acid tails (VLCFAs) [8–10]. VLCFAs that contain no double bond with 26 or more carbons in the FA acyl chain are necessary for HCMV replication [10, 11]. Saturated C16:0 FA chains are made by FA synthase, after which FA elongases (ELOVLs) can add subsequent carbons, and FA desaturases can introduce double bonds into the hydrocarbon tail. There are seven human ELOVLs, ELOVL1-7. Each ELOVL elongates FAs depending on their length and double bond content. For example, ELOVL5 elongates ≥C18 FAs that have two or more double bonds in the hydrocarbon chain (i.e., polyunsaturated FAs; PUFAs) [12, 13]. In comparison, ELOVL7 generates saturated and monounsaturated VLCFAs [10, 14, 15]. Elongated FAs can be used as substrates to support lipid synthesis.

HCMV infection increases the expression of all seven ELOVLs [8, 10, 11].

Infection shifts the host lipidome, increasing the amount of lipids that contain VLCFA tails. HCMV reprograms metabolism through several mechanisms including, but not limited to, activating sterol regulatory-element binding proteins (SREBPs) and protein kinase R (PKR)-like ER kinase (PERK), and inactivating tuberous sclerosis complex 2 (TSC2) [8, 16–20]. The process involves several viral factors, including pUL37×1, pUL38, and pUL13 [21–23]. Studies of HCMV-host metabolism interactions have focused primarily on understanding how metabolism supports virus replication, including investigations into fatty acid (FA) synthesis and elongation. Saturated VLCFAs support virus replication [10, 11], but the role of very long chain polyunsaturated FAs (VLC-PUFAs) has yet to be examined even though the synthesis of saturated VLCFAs and VLC-PUFAs are balanced in HCMV infection [8]. HCMV infection elevates the levels of lipids with VLC-PUFA tails, suggesting that they may have potential roles during virus replication [8, 23]. Moreover, some FAs may have antiviral activity [24–28].

In this study, we aimed to better understand the role of FAs in HCMV replication, either supportive or inhibitory. Consistent with previous observations, we found that HCMV infection increases the levels of most VLCFAs. However, we also noted that one monounsaturated VLCFA, C22:1 erucic acid (EA), was decreased in HCMV infected cells. EA is a key component of Lorenzo’s oil for treating X-linked adrenoleukodystrophy (X-ALD) patients [29]. Further, EA was reported to reduce influenza A virus (IAV) replication [28]. We found that EA reduces late replicative steps, causing the release of particles with reduced infectivity. EA treatment alters the lipidome of infected cells, increasing the levels of lipids with a C22:1 tail and VLC-PUFA tail. We generated ELOVL5 knockout cells (ELOVL5-KO) to determine its role in elongation of VLC-PUFAs during virus replication and its contribution to the generation of lipids containing C22:1 and VLC-PUFA tails following EA treatment. As expected, the loss of ELOVL5 caused a reduction of VLC-PUFAs and lipids that contain them in their tails. ELOVL5 does not use EA as a substrate or product in its elongation reaction; thus, it was unexpected to find that EA levels were lower in ELOVL5-depleted cells. Moreover, our experiments reveal that HCMV is more sensitive to EA treatment in ELOVL5-KO cells, suggesting that ELOVL5 promotion of lipids with C22:1 plus VLC-PUFA tails suppresses EA inhibition of HCMV. Overall, our investigation demonstrates that HCMV-induced lipid metabolism may both increase VLCFAs necessary for virus replication and reduce an antiviral.

## Results

### Erucic acid (EA) reduces HCMV replication

Some metabolites, including fatty acids (FA), can reduce virus replication in a concentration dependent manner. Here, we tested if HCMV infection affects the FA composition of a cell and whether a FA metabolite can impede HCMV replication. First, we defined FA changes that occur in response to HCMV infection by performing FA analysis using liquid-chromatography high-resolution mass spectrometry (LC-MS). We compared FAs in uninfected (mock-infected) human foreskin fibroblast cells (HFFs) to those infected with TB40/E at a multiplicity of infection (MOI) of 3 infectious unit (IU) per cell. The conditions used in our study are fully confluent, serum-starved primary fibroblasts, similar to the previous studies examining lipid metabolism following HCMV infection [8, 10, 23, 30–33]. These conditions enable us to remove any FAs or lipids that may be present in fetal bovine serum (FBS), thereby reducing the possibility that they may be measured in our assays. We identified and quantified the relative abundance of FAs ranging in length from 16 to 34 carbons (C16-C34). The relative abundance of most FAs was either increased or remained unchanged by infection (Figure 1A). The FAs that were more abundant in HCMV-infected cells than in uninfected cells were typically ≥C26 very long-chain FAs (VLCFAs), which we have previously studied [8, 10, 11, 23]. As we have previously noted, several of the FAs increased in HCMV infection were saturated or monounsaturated (e.g., C24:0, C24:1, C26:0, C26:1, C28:0, C28:1, C30:0, C30:1, C32:1, and C34:1). In contrast, the monounsaturated VLCFA C22:1 was reduced by infection (Figure 1A-B). We confirmed this observation in the lab-adapted AD169 HCMV strain (Figure 1B and S1). Both HCMV strains reduced EA by approximately 2.5-fold relative to mock-infected cells. C22:1 with a double bond at the ω9 position is called erucic acid (EA) or *cis*-13-docosenoic acid.

**Figure 1.**
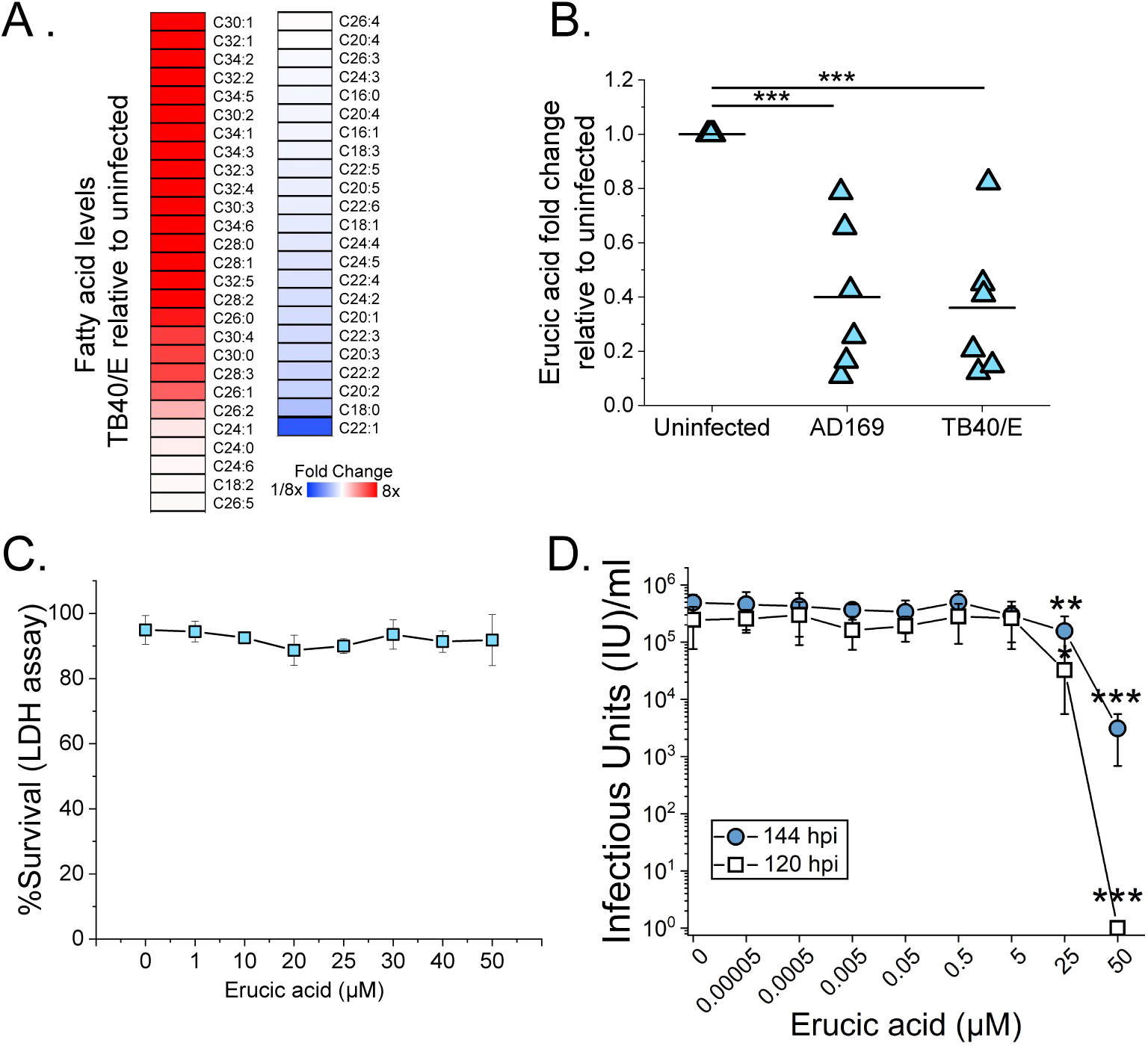
HCMV infection reduces the level of inhibitory erucic acid (EA). (A) Heatmap of fatty acid analysis in HCMV TB40/E infected HFF cells at MOI 3. The FAs were analyzed at 72hpi. (B) Quantification of EA. (C) LDH cytotoxicity assay of EA treated uninfected cells. BSA and 0.5% ethanol without EA was used as a vehicle control. (D) TCID50 assay for infectious virus production at 120 hpi and 144 hpi in HFF cells treated with EA. All data are represented as mean with standard deviation of three replicates. One-way ANOVA for (B) and Two sample t-test for (E) *p<0.05, **p<0.01, ***p<0.001

EA was reduced in the serum of two patients infected with influenza A (IAV H7N9) [34]. Moreover, EA treatment reduces IAV replication in A549 cells and mice [28]. We determine whether EA can also reduce HCMV replication. Based on the IAV observations, we hypothesized that EA treatment would reduce HCMV replication. First, we tested if EA is toxic to the cells used in our experiments by treating HFF cells with EA from 0-50 µM. We found that HFFs survived EA treatment up to 50 µM, the highest concentration we tested, suggesting that EA was not toxic to uninfected cells at 50 µM (Figure 1C). Mock-treated control cells were treated with vehicle only, 0.5% ethanol with FA-free BSA carrier. Next, we tested our hypothesis by infecting HFF cells with TB40/E at MOI 1. At 1 hpi, we washed the cells and added fresh growth medium containing 0 to 50 µM EA. At 48 hpi, the medium was replenished and supplemented with fresh EA. At 120 and 144 hpi, we collected the medium and measured the amount of infectious virus released into the supernatant. At 120 and 144 hpi, EA reduced the titers of HCMV by ∼10-fold at 25 µM (Figure 1D). EA treatment at 50 µM resulted in no measurable infectious virus at 120 hpi and reduced virus titers by ≥2.5 logs at 144 hpi (Figure 1D). Overall, we conclude that HCMV infection reduces EA levels, an endogenous FA with antiviral properties.

### EA disrupts the late stage of HCMV replication

Since EA treatment at 25 µM and 50 µM reduced HCMV titers, we investigated if EA treatment disrupted an immediate-early (IE), early (E), or late (L) replication step.

We did so by examining proteins in each of these three kinetic classes of replication. The levels of IE1 protein were unaffected by EA treatment (Figure 2A and S2A). The levels of early proteins pUL26 and pUL44 were minimally altered by EA treatment, only pUL26 was statistically reduced at 72 hpi (Figure 2A and S2A). However, we found that late viral proteins pp28 and pp71 were reduced by approximately half in EA-treated cells compared to control cells by 72 hpi (Figure 2A-B).

**Figure 2.**
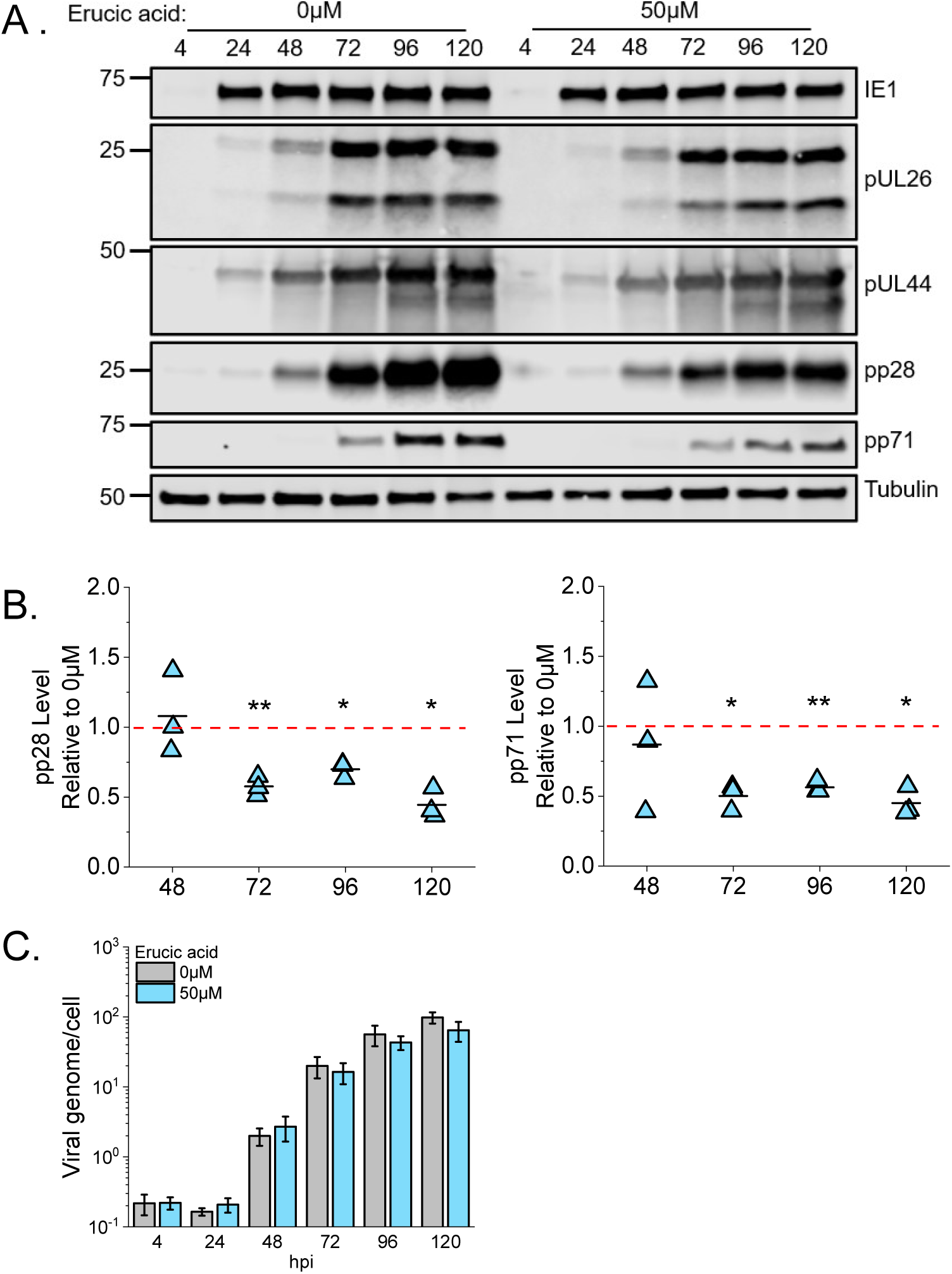
EA reduces late viral proteins. (A) Western blot of HFF infected cells treated with 50 μM EA or vehicle control (0 μM). Viral protein are immediate early (IE1), early pUL44 and pUL26, and late pp28 and pp71. (B) Quantification of late viral protein from (A). Protein levels were normalized to 0 μM of EA. Red dash line represents 1-fold. (C) Viral genomes per cell were measured using qPCR. All data are represented as mean with standard deviation. MOI 1. N=3-4. One sample t-test *p<0.05, **p<0.01, ***p<0.001

We further examined HCMV replication to determine if EA treatment disrupts an early-to-late transition step. Late gene expression depends on viral genome replication, suggesting that the reduction in late protein levels by EA treatment is due to loss in HCMV genome synthesis initiated in the early stage. We tested this possibility by determining HCMV genome levels following EA treatment. We found that the levels of viral genomes were similar in control and 50 µM EA treated cells, demonstrating that viral genome replication occurs unimpeded in the presence of EA (Figure 2C).

### EA inhibits virus replication independent of NF-κB activation

The antiviral and anti-inflammatory effects of EA in IAV infection were attributed to inactivation of NF-κB signaling [35]. NF-κB is a transcription factor that plays a key role in regulating immune responses to infection, cytokine production, cellular metabolism, and cell survival. We tested if EA inhibition of HCMV replication could also involve NF-κB signaling. To address this question, we measured NF-κB activity in the nuclei of HCMV infected cells following EA treatment. Cells were EA or vehicle-treated at 1 and 48 hpi, followed by extraction of the nuclear and cytosolic fractions at 72 hpi. We evaluated the separation of nuclear and cytosolic fractions by western blotting for tubulin (cytosolic marker) and lamin B (nuclear marker). We observed that tubulin was present in the cytoplasmic fraction while absent in the nuclear fraction (Figure S3A).

Conversely, lamin B was observed in the nuclear fraction (Figure S3A), demonstrating that our cellular fractionation protocol was successful. Next, we examined the ability of nuclear NF-κB to bind double-stranded DNA (dsDNA) using an ELISA. In this assay, NF-κB (p65) in the nuclear fraction was measured after it was bound to dsDNA sequence of the NF-κB response element. We observed that nuclear NF-κB levels were similar in cells treated with 5-50 µM EA to the level observed in untreated infected cells (Figure S3B). Our results demonstrate that EA treatment has little impact on NF-κB in HCMV-infected cells, suggesting that EA treatment inhibits HCMV replication independent of NF-κB.

### EA treatment alters the lipidome

HCMV replication depends on lipid synthesis, including FA metabolism [8, 10, 20, 23, 31]. Since EA is a FA, we determined if EA treatment disrupts the lipidome. First, we identified the effects of EA on the lipidome of uninfected cells. Here, we treated cells with 50 µM EA or mock-treated with vehicle for 72 h with EA re-feeding at 48 h to mimic the conditions used in infection. We extracted the lipids and performed lipidomics using liquid chromatography high-resolution tandem mass spectrometry (LC-MS/MS). For our analysis, we normalized lipids by cell number prior to determining the abundance of lipids in EA-treated cells relative to untreated cells. We observed that EA treatment altered the relative abundance of lipids in each of the major classes of phospholipids that were measured (Figure 3A). We found that the levels of 34 lipids (first column in the heatmap) were increased by EA treatment by 2.5-fold or greater (Figure 3A). Of the lipids increased by EA treatment, many had a sum total of 38 or more carbons among their two fatty acyl tails (Figure 3A). Moreover, we noted that these lipids typically had 1-2 double bonds among their two tails. Since EA is a C22:1 monounsaturated VLCFA, our observations suggest that EA treatment promotes the increase of lipids containing VLCFA tails with 1-2 double bonds. Additionally, we found that EA treatment reduced the relative abundance of 30 lipids by at least 40% (Figure 3A, last column). Notably, EA treatment decreased lipids with 36 or fewer carbons in their tails. Together, these findings demonstrate that a shift of the lipidome from lipids with shorter tails to longer tails occurs under EA treatment in uninfected cells. Our lipidomic findings further suggest that the exogenously fed EA molecules are entering lipid synthesis pathways to be incorporated into lipids, resulting in an increase in the number of lipids with VLCFA tails while reducing those with shorter tails.

**Figure 3.**
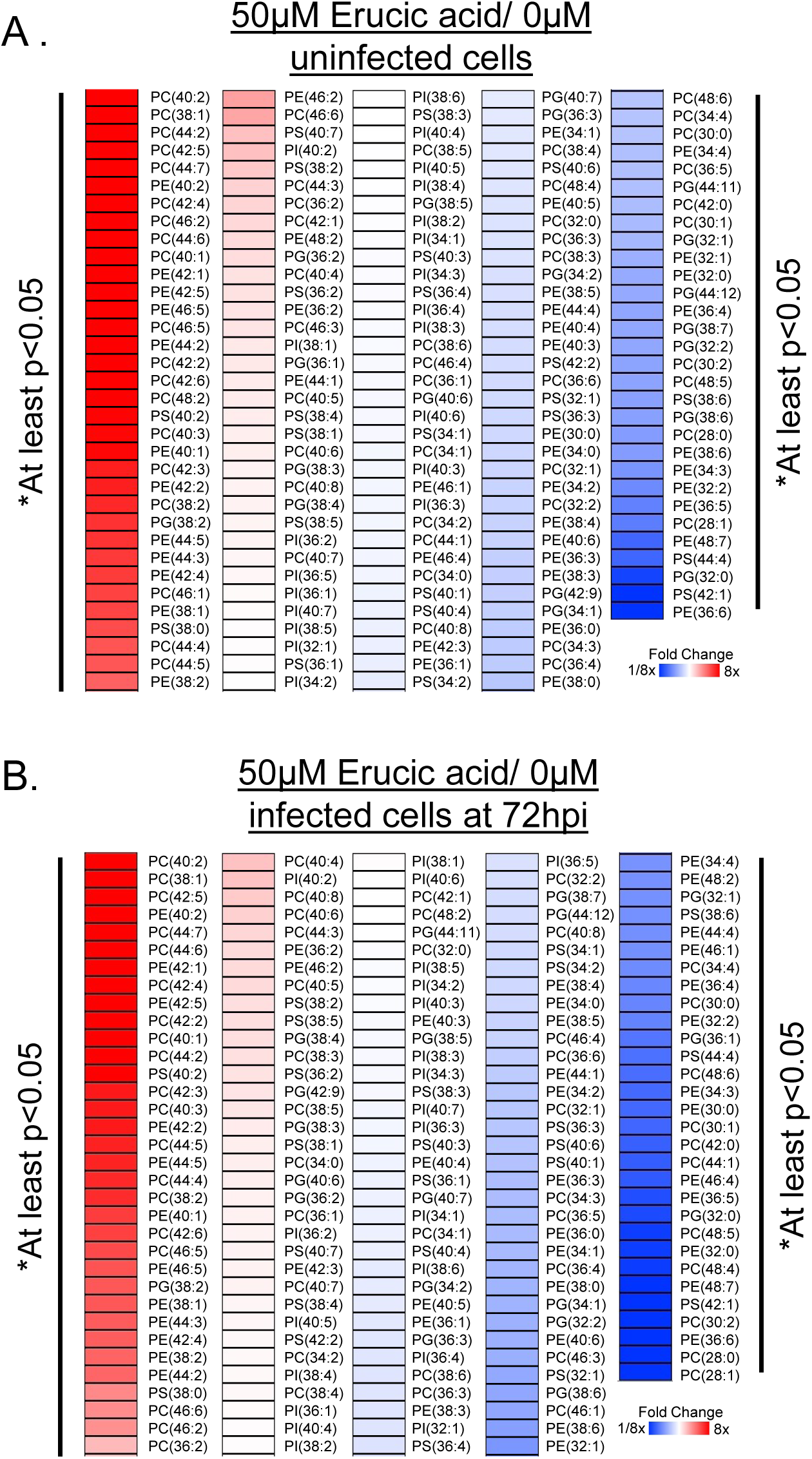
Erucic acid alters host phospholipid profile. (A) Heatmap of phospholipid analysis in EA treated uninfected HFF cells. Relative fold change of treated to untreated. (B) Phospholipid analysis of EA treated HCMV TB40/E infected cells. MOI 3. One-sample t test. N=3. *p<0.05, **p<0.01, ***p<0.001

After examining the effects of EA treatment on the lipidomic profile of uninfected cells, we determined the impact of EA treatment on the lipidome of HCMV-infected cells. As with the uninfected cells, EA treatment in HCMV-infected cells induced a shift in the host lipidome (Figure 3B). Again, EA treatment increased the relative abundance of phospholipids with tails that had a total of 38 or more carbons and 1-2 double bonds. Additionally, treatment in HCMV-infected cells resulted in lower levels of lipids with shorter tails, i.e., those that have a total ≤36 carbons among the two tails. We identified the tails of the lipids significantly reduced by EA-treatment. All of these lipids, except for three, PS(42:1), PE(48:7), and PG(38:7), lacked a C22:1 tail (Table S1). Together, these findings demonstrate that treating EA to uninfected and HCMV-infected cells generally increases the level of lipids with a C22:1 tail and reduces those lacking a C22:1 tail.

Next, we investigated the possibility that EA molecules are fed into the lipid synthesis pathway following treatment by identifying the tails of the lipids increased most by EA treatment. The tail identification was done using MS/MS fragmentation of the lipids that were increased by ≥2-fold in EA treatment. Of these 34 lipids, we observed a C22:1 tail in 31 of them (Table 1). This was the case in both uninfected and HCMV-infected cells and provides additional support that treatment promotes an increase in the level of lipids containing an EA tail.

**Table 1.** Fatty acid composition in lipids that were upregulated by EA treatment.

| Lipid | Observed Tails | Fold Difference relative to uninfected 0 μM EA | | | |
| --- | --- | --- | --- | --- | --- |
|  |  | Uninfected |  | Infected |  |
| | | 0 μM | 50 μM | 0 μM | 50 μM |
| PC(40:2) | C22:1, C18:1 | 1 | 106.9 | 1.0 | 97.8 |
| PC(38:1) | C22:1, C16:0, C18:1, C20:1 | 1 | 55.6 | 0.9 | 32.2 |
| PC(44:2) | C22:1, C18:1, C26:1 | 1 | 50.3 | 3.0 | 30.1 |
| PC(42:5) | C22:1, **C20:4** | 1 | 44.6 | 0.7 | 20.1 |
| PC(44:7) | C22:1, C18:0, **C22:6** | 1 | 35.1 | 1.1 | 23.0 |
| PE(40:2) | C22:1, C18:1, C16:1 | 1 | 24.9 | 1.0 | 23.8 |
| PC(42:4) | C22:1, C18:1, **C20:3, C20:4, C22:3** | 1 | 22.4 | 0.8 | 14.0 |
| PC(46:2) | C22:1, C18:1, C24:1, C28:1 | 1 | 20.9 | 9.8 | 23.0 |
| PC(44:6) | C22:1, C18:1, **C20:5, C22:5**, C24:1, | 1 | 18.8 | 0.7 | 13.8 |
| PC(40:1) | C22:1, C16:0, C16:1, C18:1, C24:1 | 1 | 16.5 | 1.0 | 10.2 |
| PE(42:1) | C22:1, C16:1, C18:0, C18:1, C24:0, C24:1 | 1 | 16.1 | 0.6 | 11.2 |
| PE(42:5) | C22:1, C18:1, **C20:3, C20:4, C22:4, C24:5** | 1 | 14.2 | 1.2 | 16.0 |
| PE(46:5) | **C20:4, C22:4, C22:5**, C24:0, C26:1 | 1 | 13.1 | 2.3 | 8.5 |
| PC(46:5) | C22:1, **C20:3, C20:4, C22:5**, C26:1 | 1 | 12.9 | 2.0 | 7.7 |
| PI(40:2) | C22:1, C18:1 | 1 | 9.7 | 2.4 | 24.4 |
| PE(44:2) | C22:1, C18:1, C20:1, C24:1, C26:1 | 1 | 9.4 | 1.7 | 5.5 |
| PC(42:2) | C22:1, C16:1, C18:1, C20:1, C24:1, C26:1 | 1 | 9.0 | 1.2 | 13.6 |
| PC(42:6) | C22:1, C18:1, **C20:5, C22:5, C24:5** | 1 | 8.4 | 0.9 | 4.2 |
| PC(48:2) | C22:1, C18:1, C24:1, C26:1, C30:1 | 1 | 8.0 | 27.8 | 24.8 |
| PS(40:2) | C22:1, C16:0, C18:1 | 1 | 7.8 | 0.7 | 6.1 |
| PC(40:3) | C22:1, C16:0, C18:1, **C18:2, C20:2, C22:2, C22:3, C24:3** | 1 | 7.7 | 0.9 | 5.5 |
| PE(40:1) | C22:1, C18:0, C20:1, C22:0, C24:0 | 1 | 7.6 | 1.0 | 4.8 |
| PC(42:3) | C22:1, C16:1, C18:0, C18:1, **C18:2, C20:3, C20:4**, C24:1 | 1 | 6.3 | 0.9 | 5.8 |
| PE(42:2) | C22:1, C18:1, C18:2, C20:1, C24:0, C24:1 | 1 | 6.3 | 1.0 | 6.1 |
| PC(38:2) | C22:1, C16:1, C18:1, C20:1, **C20:2** | 1 | 5.8 | 1.2 | 6.4 |
| PG(38:2) | C22:1, C16:1, C18:1, C20:1, **C20:2** | 1 | 5.3 | 1.7 | 6.8 |
| PE(44:5) | C20:4, C24:1 | 1 | 5.3 | 1.0 | 6.0 |
| PE(44:3) | C22:1, C18:1, **C18:2, C20:2, C20:3, C22:2**, C24:1 | 1 | 5.1 | 1.1 | 3.9 |
| PC(46:1) | C22:1, C16:1, C18:1, C24:0, C28:0, C30:1 | 1 | 4.7 | 11.8 | 4.4 |
| PE(38:1) | C22:1, C18:0, C18:1, C20:1, C22:0 | 1 | 4.6 | 1.2 | 4.5 |
| PS(38:0) | C16:0, C18:0, C20:0 | 1 | 4.3 | 1.7 | 4.7 |
| PC(44:4) | C22:1, C18:1, **C18:2, C20:3, C20:4, C22:3**, C24:1, **C26:3** | 1 | 4.0 | 1.0 | 5.3 |
| PC(44:5) | C22:1, C18:1, **C20:4, C22:4**, C24:1 | 1 | 4.0 | 0.8 | 4.4 |
| PE(38:2) | C22:1, C16:1, C18:0, C18:1, **C18:2, C20:2**, C20:1 | 1 | 3.6 | 1.5 | 5.2 |

Polyunsaturated fatty acids are shown in bold.

In addition to the alterations in the cellular lipidome described above, we also noted that EA-treatment impacted the levels of several lipids with polyunsaturated (PUFA) tails (Table 1). PUFAs contain two or more double bonds in the hydrocarbon chain. While some lipids with a total of 3-4 double bonds were increased by EA-treatment, we also found that lipids with 5-7 double bonds were elevated in uninfected and HCMV-infected cells (Figure 3 and Table 1). Almost half of the lipids significantly increased by EA-treatment contained a PUFA tail (Table 1). Moreover, 15 of the 31 lipids that contained an EA tail also had a PUFA tail, suggesting that the synthesis of PUFAs may also promote the incorporation of EA into lipids.

### HCMV infection elevates fatty acid elongase 5 (ELOVL5)

Since we previously demonstrated that lipids with saturated or monounsaturated VLCFAs are increased by HCMV infection [8, 10, 23], we decided to focus on the observation that some lipids with an EA tail also have a very-long chain PUFA (VLC-PUFA) tail (Table 1). Prior to attachment to a lipid backbone, shorter chain PUFAs are elongated by fatty acid elongases (ELOVLs). ELOVL2, 4, and 5 elongate PUFAs to produce VLC-PUFAs. A screen of all seven ELOVLs suggested that ELOVL5 may be important to HCMV infection [10] and we previously demonstrated that HCMV infection increases ELOVL5 levels [8]. Thus, we investigated if there is a relationship between EA and ELOVL5 regarding HCMV replication and lipidome remodeling. However, before we can better understand if ELOVL5 impacts PUFA metabolism in EA treatment and its antiviral properties, we must determine if ELOVL5 plays a role in HCMV replication.

While it is known that ELOVL5 protein levels are increased during the late stages of HCMV replication [8], the levels of ELOVL5 protein during earlier stages of virus replication have not been evaluated. We measured the protein level of ELOVL5 from 4 to 144 hpi after infecting fibroblast cells at MOI 1 with TB40/E-GFP. ELOVL5 protein levels increased by 48 hpi and remained elevated through 144 hpi (Figure 4A).

**Figure 4.**
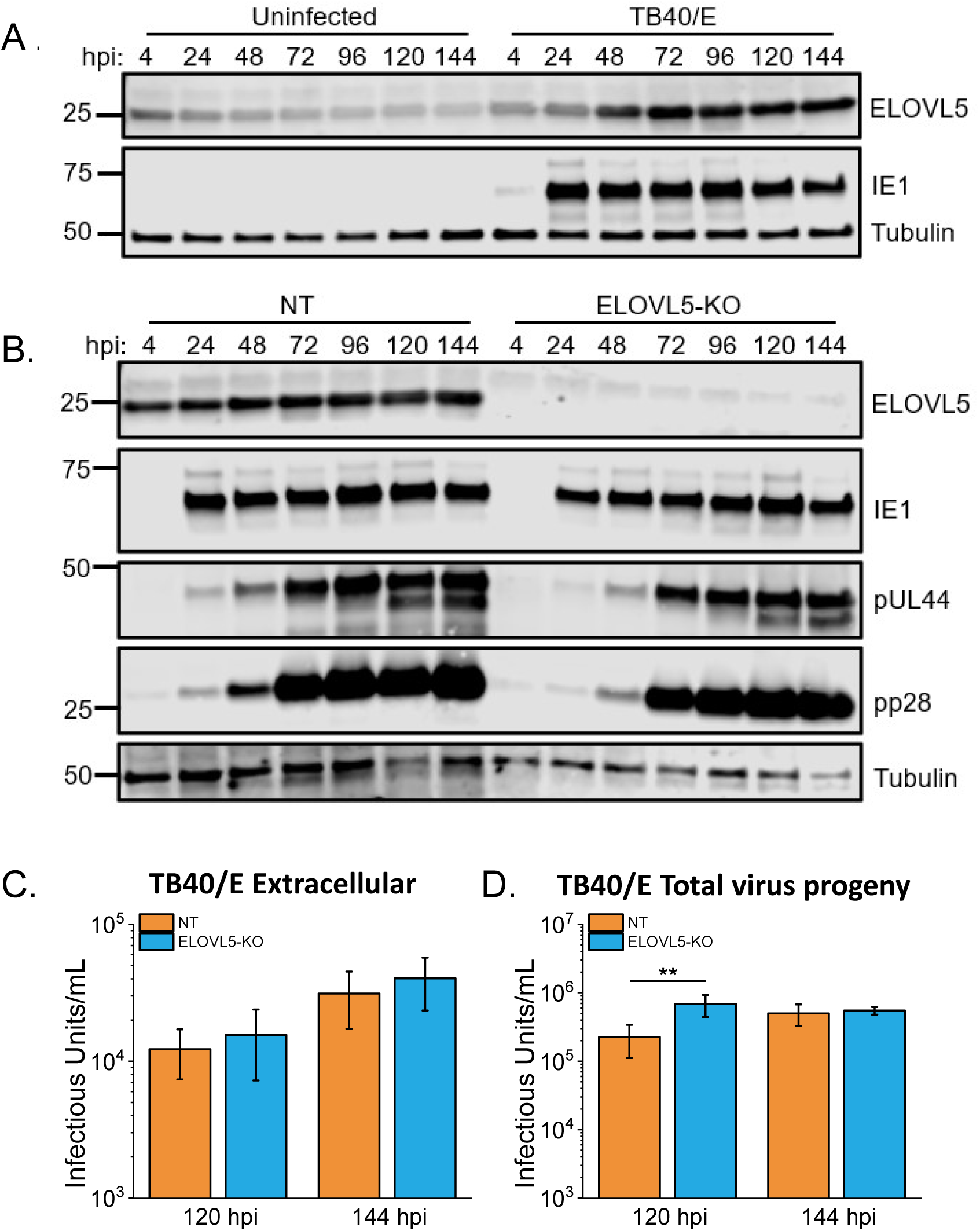
HCMV replication does not require ELOVL5. (A) ELOVL5 protein levels were measured by western blot during HCMV replication. (B) Western blot ELOVL5 CRISPR/Cas9 knockout (ELOVL5-KO) cells and non-targeting (NT) control cells. (C) The release of infectious progeny in the supernatant was measured by TCID50. (D) Total virus progeny (cell-free and cell associated) was measured by TCID50. MOI 1. Two sample t-test. **p<0.01

Since infection promotes ELOVL5 levels, we investigated if it supports HCMV replication. We generated ELOVL5 knockout (ELOVL5-KO) cells using CRISPR/Cas9. The loss of ELOVL5 was evaluated by western blotting (Figure 4B) and confirmed by sequencing (Figure S4A). In parallel, we generated control fibroblasts that express a non-targeting (NT) gRNA that lacks the ability to target any human or HCMV gene. With these CRISPR/Cas9 cells, we investigated if the loss of ELOVL5 affected the levels of HCMV proteins. The levels of immediate-early (IE1), early (pUL44), and late (pp28) proteins were similar in ELOVL5-KO and NT control cells (Figure 4B). To further investigate if ELOVL5 provides a function that supports HCMV replication, we used these cells to determine if ELOVL5 is required for the production of HCMV infectious progeny. We measured the number of infectious TB40/E virions in the extracellular medium at 120 and 144 hpi following infection at MOI 1. At both times, a similar number of infectious progeny was produced by the ELOVL5-KO and NT control cells (Figure 4C). Since TB40/E infectious virus is both cell-associated and extracellularly released, we measured total virus production (cell-associated plus released virions). At 120 hpi, the production of total infectious progeny was statistically significantly increased in ELOVL5-KO cells relative to NT cells, but the increase was only 3-fold (Figure 4D).

However, no difference in virus titers between ELOVL5-KO and NT cells was observed at 144 hpi (Figure 4D). We further evaluated if ELOVL5 contributes to the replication of HCMV by evaluating its impact on the level of infectious virus produced by cells infected with the lab-adapted AD169 strain. We infected NT and ELOVL5-KO cells using AD169 at MOI 0.5 and 1 and measured extracellularly released infectious virus progeny at 96-144 hpi (Figure S4B-C). Again, ELOVL5-KO and NT control cells produced similar amounts of infectious HCMV progeny, except at 120 hpi in cells infected at MOI 1, where the titers from ELOVL5-KO were 3.5-fold lower than NT control cells (Figure S4C). Taken together, these results indicate that ELOVL5 is likely dispensable for HCMV replication under these conditions.

### ELOVL5 is active in HCMV-infected cells

Since ELOVL5 is unnecessary for HCMV replication, it may not be active in infected cells even though the protein levels increase. To test if ELOVL5 is active and contributes to FA elongation in the conditions used in this study, we performed a FA analysis in ELOVL5-KO and NT cells. ELOVL5 produces VLC-PUFAs that are typically ≥20 carbons in length using C18:2-C18:4 as substrates [12, 13, 36]. As we did earlier, we measured FAs using LC-MS to gain insight into the impact of ELOVL5 on the FA profile of uninfected and infected cells. The level of C18:3 was significantly higher in uninfected ELOVL5-KO cells compared to NT cells (Figure 5A). These results suggest that C18:3 is accumulating due to a reduction in their elongation to longer FAs in ELOVL5-KO cells. The levels of several other FAs, including C18:2, were elevated; however, their increase was not statistically significant. Additionally, the loss of ELOVL5 resulted in a significant decrease in several FAs, including the expected products:

**Figure 5.**
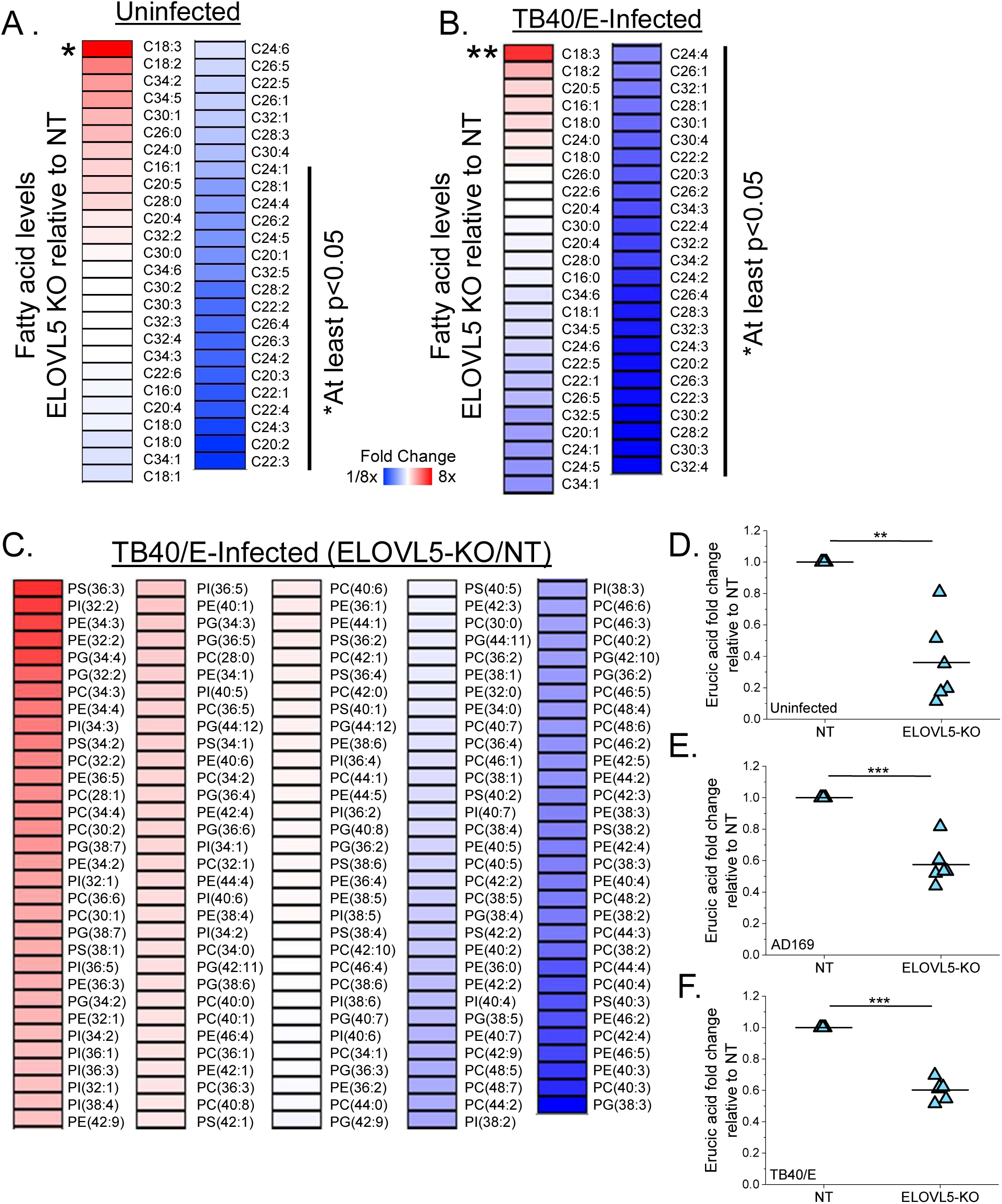
Depletion of ELOVL5 reduces VLC-PUFAs and EA. (A) FA analysis in uninfected cells. Relative fold change of ELOVL5-KO to NT cells. (B) FA analysis in TB40/E-infected cells. (C) Lipid profile in HCMV-infected ELOVL5-KO cells relative to infected NT control cells. (D-E) Relative level of EA in uninfected (D), AD169-infected (E), and TB40/E-infected (F) cells. (B-F) MOI 3. 72 hpi. (A-F) One sample t-test. *p>0.05, **p<0.01, ***p<0.001

C20:2-C20:3, C22:2-C22:4, and C24:2-C24:3 (Figure 5A). These findings are consistent with ELOVL5 producing VLC-PUFAs in uninfected fibroblasts from shorter chained desaturated FAs [12, 13, 36]. Next, we examined the role of ELOVL5 in FA elongation in HCMV-infected cells. The total FA abundance in HCMV TB40/E or AD169 infected ELOVL5-KO cells was compared to infected NT cells. Again, C18:3 levels were higher in the infected ELOVL5-KO cells, further indicating that it is a substrate of ELOVL5 (Figure 5B and S5). The loss of ELOVL5 reduced the relative abundance of several FAs in HCMV-infected cells, including VLC-PUFAs, similar to uninfected cells.

We further investigated the activity of ELOVL5 in HCMV infection by analyzing the lipid content in ELOVL5-KO cells compared to NT control cells. At 72 hpi, we observed that 26 lipids were higher by ≥2-fold when ELOVL5 was depleted (Figure 5C). Half of these 26 lipids have three or more double bonds among the two FA tails. We further analyzed these lipids by identifying their tail composition using MS2 and found that several contained a C18:3 tail in the ELOVL5-KO (Table S2). This observation is consistent with an increase in the pool of ELOVL5 substrates, primarily C18:3, that can be used to support lipid synthesis.

Additionally, we also observed lipids downregulated upon knockout of ELOVL5 in HCMV-infected cells (Figure 5C). We found that 32 lipids were significantly decreased in ELOVL5-KO relative to NT control cells. Identification of the tails revealed that many have a ≥C22 VLC-PUFA tail that contained two or more double bonds (Table S3).

Further, we were able to collect MS2 information on some lipids in the ELOVL5-KO samples (e.g., PC(42:4) in Table S3). This may be due to their low abundance in the absence of ELOVL5. Overall, we conclude that ELOVL5 is active in uninfected and HCMV-infected fibroblasts and is elongating C18:3 to generate longer PUFAs.

### ELOVL5 affects the levels of EA, a monounsaturated C22:1 FA

While most of the FAs reduced in ELOVL5-KO cells were VLC-PUFAs, we noted that some were monosaturated—those with one double bond (Figure 5A-B). Notably, C22:1 is lower in uninfected ELOVL5-KO cells compared to NT cells (Figure 5D). In AD169 and TB40/E infection, EA is also reduced in ELOVL5-KO cells (Figure 5E-F).

The observation that ELOVL5 may regulate the levels of EA was unexpected since C22:1 is not known to be a substrate or product of ELOVL5. These findings suggest that ELOVL5 may have a role in regulating EA levels.

### ELOVL5 supports the incorporation of EA into lipids

Since ELOVL5 may potentiate regulation of EA levels and EA treatment increases lipids with a C22:1 plus VLC-PUFA tails (Figures 3 and 5), we investigated if ELOVL5 contributes to the shift in the lipidome caused by EA treatment. We performed lipidomics in ELOVL5-KO cells and NT following EA or vehicle treatment. For uninfected cells, we treated the ELOVL5-KO cells with 50 µM of EA or vehicle at 1 h post mock-infection, replenished the media with EA at 48 h post treatment, and extracted lipids at 72 h post initial treatment. Similar to our previous observation, EA treatment in ELOVL5-KO cells caused a broad shift in the lipidome of uninfected cells (Figure 6A). The same observation was made in HCMV-infected cells (Figure 6B). In this case, the lipids increased the greatest were typically those that contained a C18:3 tail that were elevated in untreated ELOVLs-KO cells (e.g., PC(36:3) in Figure 5C and Table S2). Further, we observed that some lipids containing a C22:1 tail, which increased in non-CRISPR/Cas9 cells following EA treatment, also increased in treated ELOVL5-KO cells. However, their level of increase was reduced in ELOVL5-KO cells (e.g., PC(40:2) in Figure 3 and Table 1). This was also the case for lipids that contained a combination of a C22:1 tail plus a VLC-PUFA tail (e.g., PE(42:5)). We conclude that EA is likely incorporated into lipids but does so at a lower level in the absence of ELOVL5. These observations suggest that ELOVL5, through its activity of elongating VLC-PUFAs, helps support EA metabolism into lipids.

**Figure 6.**
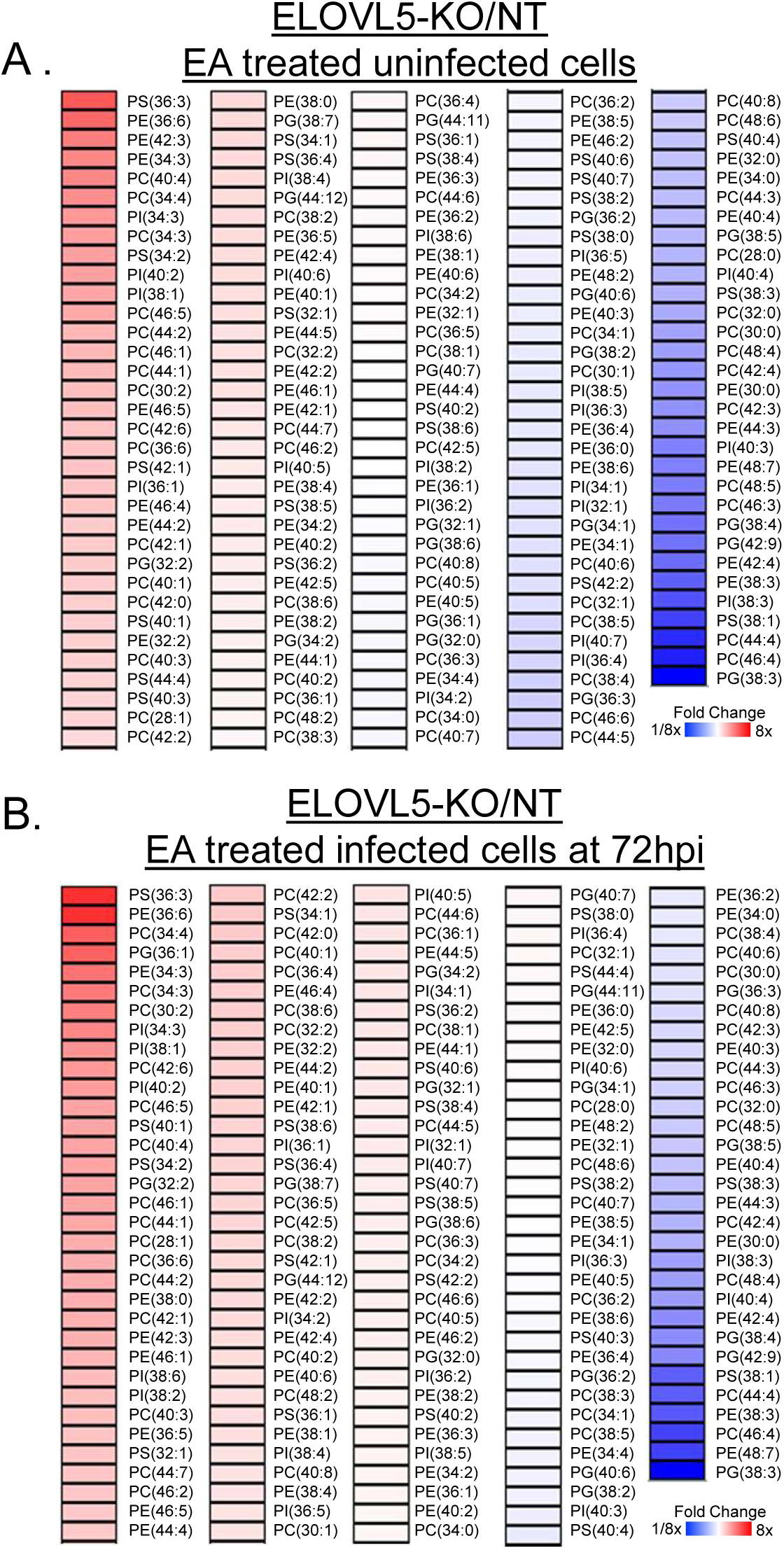
ELOVL5 enhances the levels of EA containing lipids. (A) Phospholipid levels in 50 µM EA treated uninfected ELOVL5-KO cells. Relative fold change of treated ELOVL5-KO to treated NT cells. (B) Phospholipid analysis in 50 µM EA treated HCMV infected ELOVL5-KO cells. Relative fold change of infected treated ELOVL5-KO to infected treated NT cells. MOI 3, 72 hpi. One-sample t test. N=3. *p<0.05, **p<0.01, ***p<0.001

### ELOVL5 diminishes EA inhibition of HCMV replication

Since ELOVL5 may regulate EA metabolism, we examined if it affects EA inhibition of HCMV replication. Since EA treatment reduced the levels of late viral proteins, we started by examining the effect of EA on viral proteins in ELOVL5 depleted cells. IE1 protein levels were similar in NT and ELOVL5-KO cells under 50 µM EA-treatment (Figure 7A). Early proteins pUL26 and pUL44 were slightly reduced in EA-treated ELOVL5-KO cells. Notably, late proteins pp28 and pp71 were reduced at late times of replication in EA-treated ELOVL5-KO cells (Figure 7A-B). Since late proteins were reduced, we measured viral genome replication. The levels of viral genomes were similar in ELOVL5-KO and NT cells treated with 50 µM EA (Figure 7C), suggesting EA treatment disrupts late replication and may impact early stages independent of viral genome replication in the absence of ELOVL5.

**Figure 7.**
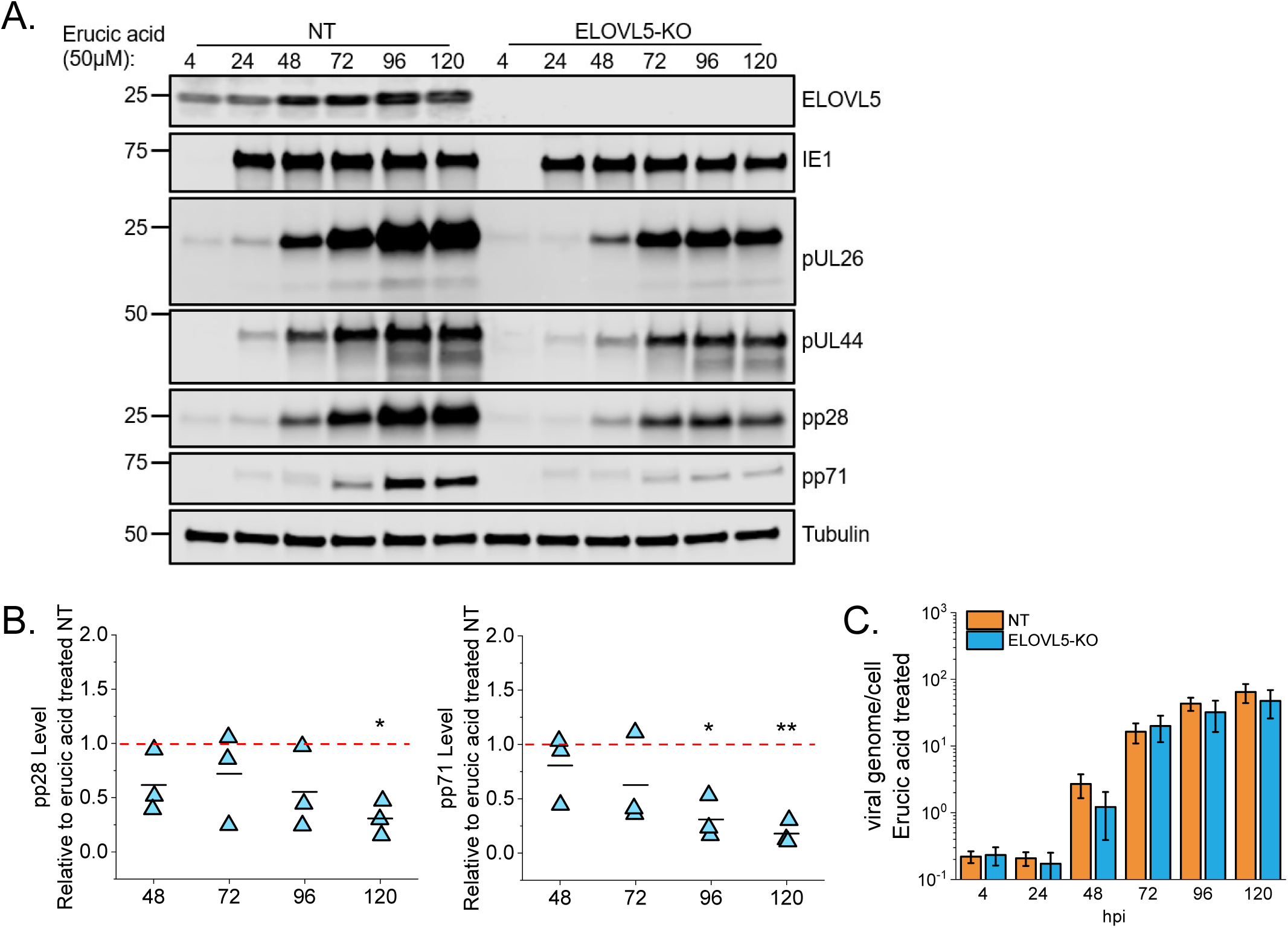
ELOVL5 diminishes EA suppression of late replication. (A) Western blot of viral protein levels in 50 µM EA treated ELOVL5-KO cells compared to NT cells. (B) Quantification of late viral protein levels in A. (C) Viral genomes per cell in 50 µM EA treated infected ELOVL5-KO cells compared to NT cells. N=3-4. One sample t-test *p<0.05, **p<0.01, ***p<0.001

Since our measurements of viral proteins suggest that EA treatment disrupts HCMV replication to a greater extent in ELOVL5-KO cells, we investigated whether ELOVL5 regulates the impact of EA on virus titers. For NT cells, EA treatment reduced the amount of infectious progeny released into the supernatant at 25 µM and 50 µM (Figure 8). This was consistent with our earlier observation in non-CRISPR/Cas9 treated cells (Figure 1). In ELOVL5-KO cells, EA treatment further decreased infectious virus production by 3-fold at 25 μM and more than 100-fold at 50 μM EA relative to the NT cells (Figure 8). This suggests that the loss of ELOVL5 makes HCMV more sensitive to the antiviral properties of EA. Overall, the reduction of late viral proteins and the loss of progeny production in the EA-treated ELOVL5-KO cells suggest that ELOVL5 ameliorates the effects of EA on HCMV replication.

**Figure 8.**
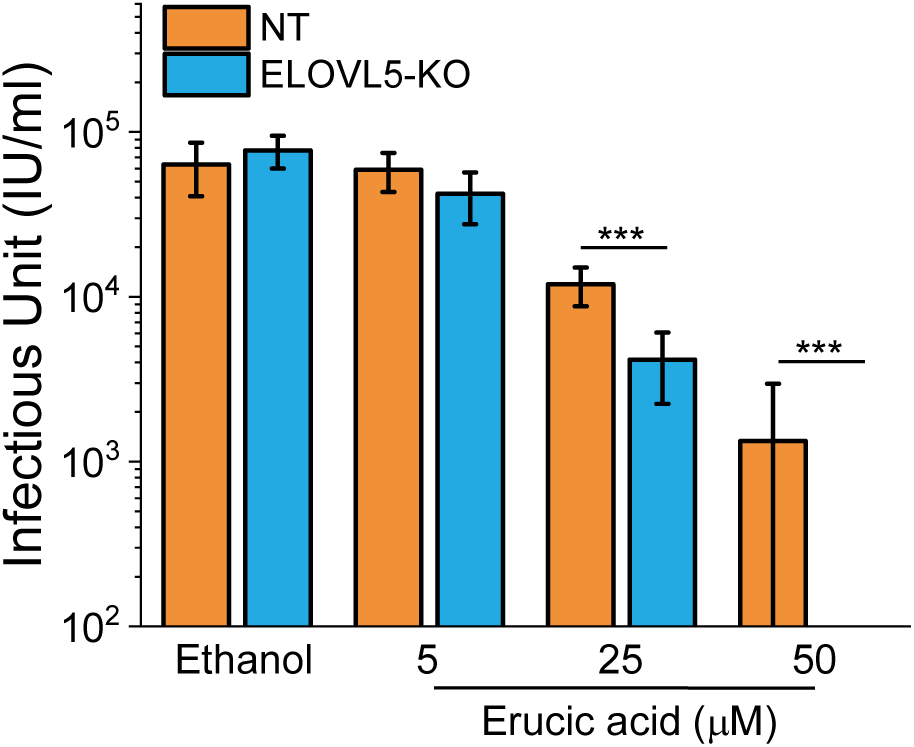
ELOVL5 mollifies EA inhibition of HCMV. ELOVL5-knockout and NT cells were EA or vehicle treated at 1 hpi. The amount of infectious progeny released by cells was measured by TCID_50_. Some samples were below the limit of quantification in the 50 µM EA treated samples. Two sample t-test. MOI 1, 120 hpi. N=3. Two sample t-test. ***p<0.001

### EA reduces HCMV infectivity, ELOVL5 lessens its impact on infectivity

Since EA treatment disrupts late steps in virus replication, we measured its impact on virus particle release. Virus particles released in the extracellular environment were measured by quantifying the number of enveloped, encapsidated HCMV genomes released into the supernatant. In this case, cells infected at MOI 1 were treated with 50 µM EA at 1 and 48 hpi, followed by particle measurements using qPCR. EA treated and vehicle treated cells released similar amounts of virus particles, suggesting that EA likely has little to no effect on HCMV particle production or their egress (Figure 9A). This was also the case in ELOVL5-KO and NT cells (Figure 9B).

**Figure 9.**
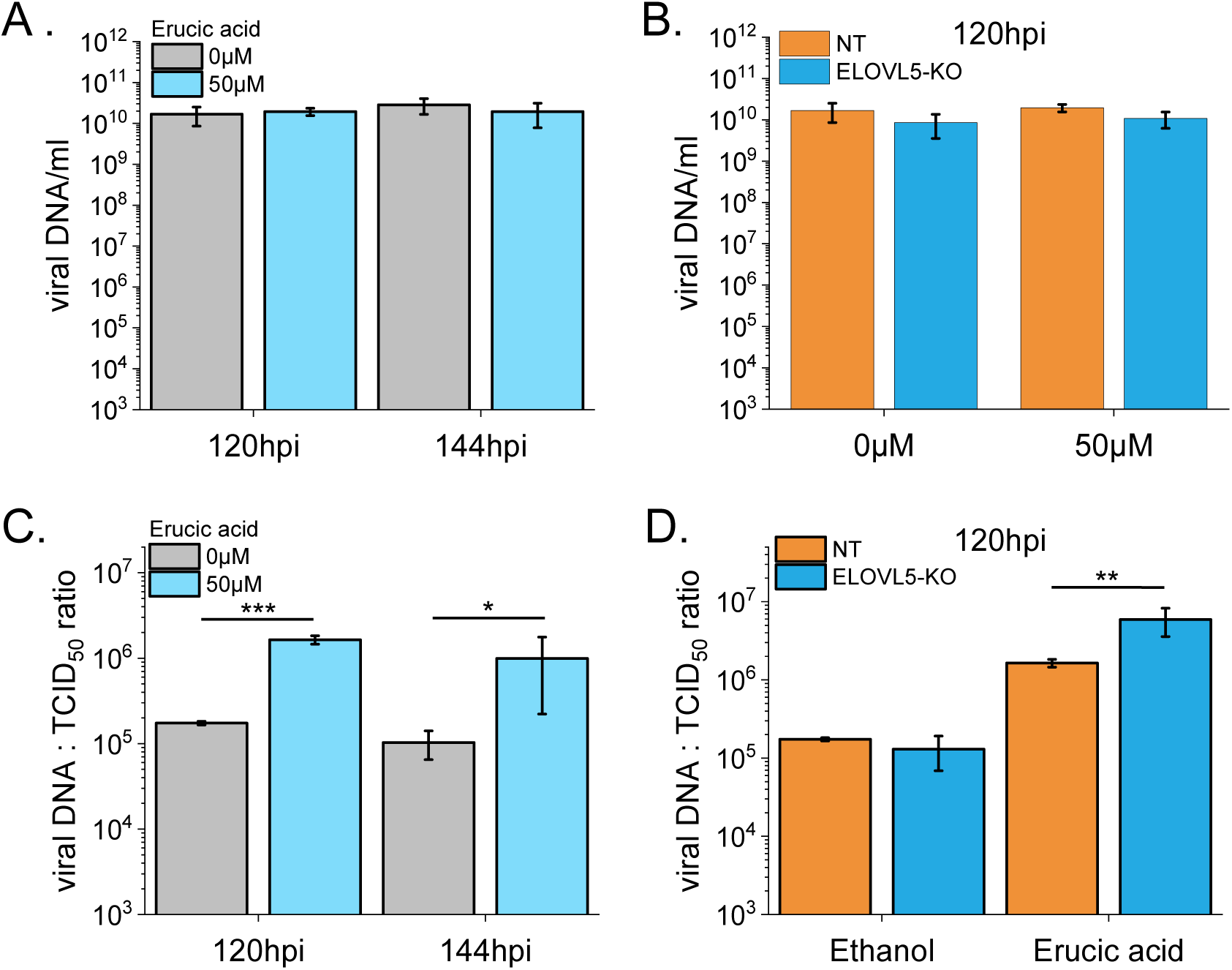
EA decreases HCMV infectivity and ELOVL5 lessens the effect of EA. (A) The amount of released virus particles was measured in the supernant by qPCR in EA treated and untreated HFF cells. (B) Released virus particles in treated and untreated NT and ELOVL5-KO cells. (C) The ratio of released total particles to infectious particles was compared for treated and untreated HFF cells shown in A. (D) The ratio of released total particles to infectious particles was compared for treated and untreated NT and ELOVL5-KO cells shown in B. MOI 1. N=3. Two sample t-test in log-transformed value. *p<0.05, **p<0.01

Given that EA-treatment can reduce HCMV titers by several logs but has little to no effect on the number of particles released, we determined if the virus particles released by EA-treated cells retain their ability to be infectious. We determined the infectivity of the released particles by comparing the ratio of the total number of released particles to those that are infectious (i.e., the particle-to-infectious virion ratio). Cells treated with 50 µM EA released fewer infectious virions per total number of released particles when compared to vehicle treated cells (Figure 9C). EA treatment caused a 10-fold loss in infectivity in these cells. Next, we investigated if ELOVL5 impacts EA’s effects on infectivity. In the vehicle treated cells, the infectivity of viruses produced by NT and ELOVL5-KO cells was similar, demonstrating that ELOVL5 is not required for HCMV infectivity (Figure 9D). While NT cells treated with 50 µM EA released a similar number of particles as untreated NT cells, fewer of those particles were infectious, resulting in an approximately 10-fold reduction in infectivity (Figure 9B). We conclude that ELOVL5 limits the ability of EA to reduce HCMV replication, including the infectivity of released virus particles.

## Discussion

All viruses rely on host machinery to obtain energy and building blocks from the cells for successful replication. For HCMV, this involves promoting FA synthesis and elongation that is necessary for virus replication [10, 11, 31]. Viral infection triggers host metabolic reprogramming in the cells, which provides free nucleotides for viral genome replication and increased amino acids for the building blocks of viral proteins. Many viruses alter central carbon metabolism, including glycolysis, glutaminolysis, TCA cycle, nucleotide synthesis, amino acid metabolism, as well as fatty acid and lipid synthesis [37–42]. Alterations in metabolism during virus infection can promote the synthesis of metabolites and FAs with antiviral properties [43, 44]. Consequently, viruses may have evolved mechanisms to ensure that metabolic products required for their success are made while those that may inhibit replication have minimal impact on the virus.

The increase in fatty acid synthesis and elongation caused by HCMV infection leads to a shift toward VLCFAs [10, 11, 23, 31]. As we previously noted, HCMV infection increases the levels of saturated and monounsaturated VLCFAs (Figure 1 and S1) [8, 10, 11, 23]. HCMV shifts the lipidome by promoting lipid synthesis via inducing multiple lipogenic genes involved in fatty acid and lipid synthesis pathways, including ELOVLs [9, 10, 16, 18, 45, 46]. HCMV increases ELOVL7 and requires its activity for replication [9, 10, 46]. Infection also increases ELOVL5 expression and protein levels (Figure 4A) [10, 46]. The increase in ELOVL7 in HCMV infection requires the ER-stress response kinase, PERK; however, the increase in ELOVL5 is PERK-independent [8].

While all ELOVLs are regulated, at least in part, by sterol regulatory element-binding proteins (SREBPs), our previous study in HCMV demonstrated that the levels of ELOVL are differentially regulated in ways that are not well understood [8]. In this study, we demonstrated that ELOVL5 is active during HCMV replication (Figure 5 and S5). In HCMV infected cells, ELOVL5 elongates C18:3, and possibly C18:2, to generate VLC-PUFAs (Figure 5 and S5). Consistent with this observation, we found that infection increases the levels of VLC-PUFAs with at least 2-3 double bonds (Figure 1A and S1). A previous shRNA knockdown screen in MRC-5 cells growing in FBS-containing media suggested that ELOVL5 was necessary for virus replication [10]. We found under serum-free growth conditions used in this study that ELOVL5 is not required for HCMV replication in HFF cells (Figures 4 and S4). ELOVL5 may support HCMV replication in MRC-5 but not HFFs, or differences in the conditions may alter the necessity of ELOVL5, including the potential FAs present in the experiments. Nonetheless, our study demonstrates that ELOVL5 supports HCMV replication under conditions where the cells are fed an exogenous FA (Figure 8-9).

In contrast to HCMV elevation of monounsaturated VLCFAs, infection decreased the C22:1 monounsaturated VLCFA EA (Figure 1A-B and S1). We found that EA suppresses HCMV replication at concentrations greater than 25 µM, including those that are non-cytotoxic to HFF cells (Figure 1C-D). EA suppressed IAV replication, potentially by targeting NF-κB and p38 MAPK activation [28]. Notably, EA did not activate NF-κB in HCMV infected cells (Figure S3). Instead, EA treatment disrupted the late stage of HCMV replication, resulting in the release of particles with a loss of infectivity (Figures 1-2, 9). The disruption could be due to an indirect effect of EA. Alternatively, EA may directly target EA replication by interacting with viral proteins or the lipids of the virus envelope. Notably, we demonstrate that EA’s ability to suppress HCMV is partially mollified by ELOVL5 (Figures 8-9). Moreover, we found that EA levels were reduced in the absence of ELOVL5 in uninfected and infected cells (Figure 5). Our observations suggest that ELOVL5 regulates EA levels through a mechanism that does not involve the enzyme acting directly on EA. ELOVL1 can produce and elongate EA [12]. HCMV increases the expression of ELOVL1; however, its potential role in virus replication remains unknown [10].

ELOVLs have a preference for substrates and products that depend on the chain length and double bond content. In addition to ELOVL5, VLC-PUFAs are made by ELOVL2. HCMV infection induces ELOVL2 expression [10]. Loss of ELOVL5 in mouse livers may result in a compensatory effect leading to increased activity of ELOVL2 [13]. The ELOVL5-KO mice developed hepatic steatosis due to increased triacylglycerol levels caused by activation of SREBPs, which promotes ELOVL2 expression and several other lipogenic genes, including glucose transporter type 4 (GLUT4), acetyl-CoA carboxylase (ACC), and FA synthase [13, 18, 30, 31]. In ELOVL5-WT infected cells, HCMV infection increases the expression and activity of GLUT4, ACC, and FA synthase [47]. A feedback regulation or any other possible relationship between ELOVL5 and SREBP activities in HCMV infection is unknown; however, the ELOVL5-KO mouse findings suggest a positive-negative feedback loop between ELOVL5 and SREBP1c via the levels of VLC-PUFAs [36]. While ELOVL2 was shown to be dispensable for HCMV replication [10], it may provide an essential function in ELOVL5-KO cells or assist ELOVL5 activity in EA-treated cells. Even so, we found that ELOVL5 plays a role in HCMV replication under EA treatment.

In addition to HCMV, ELOVLs have been studied in a few other viruses.

ELOVL5, along with ELOVL4 and 7, are increased by hepatitis C virus (HCV) infection [48]. Even though PUFA levels are increased in HCV-infected cells [49, 50], ELOVL5 knockdown did not affect HCV replication [48], and its overexpression only slightly enhances virus replication [51]. Hepatitis A virus (HAV) increases the levels of lipids with VLC-PUFA tails, primarily in sphingolipids [52]. In addition to lipids with VLC-PUFA tails, HAV infection also increased lipids with a C22:1 tail. HAV infection increased the expression of ELOVL4 and 7; however, ELOVL5 expression remained unchanged [52]. The synthesis of VLCFAs aided HAV genome replication, and sphingolipids supported the assembly and release of HAV quasi-enveloped virions [52]. Overall, HCV and HAV, along with our HCMV findings, demonstrate that diverse viruses may promote different ELOVLs and increase lipids with VLC-PUFAs to support various steps in virus replication.

Based on our findings, we propose a model for ELOVL5 regulation of EA and suppression of its ability to inhibit HCMV (Figure 10A). While we demonstrate that EA reduces late protein levels and causes a loss in infectivity, the antiviral mechanism of action remains undefined. We demonstrate that EA treatment does not alter NF-κB activity, suggesting the antiviral mechanism is independent of NF-κB and p38 MAPK signaling. A previous study in IAV infection showed that EA suppressed NF-κB and p38 MAPK activity; however, they were treating EA at ≥300 µM, whereas we found inhibition of HCMV at 25-50 µM [28]. EA is a C22:1 VLC-monounsaturated FA, similar long-chain or VLC monounsaturated FAs are known to have virus inhibitory properties. Treatment of C18:1 and C18:2 blocks several enveloped DNA and RNA viruses, including HSV-1, HSV-2, IAV, Sendai virus, and West Nile virus [25–27]. The FA likely reduces an early step in replication [26, 27]. Direct incubation of the FA with the virus resulted in a loss of infectivity due to disruption or partial lysis of the virus membrane [27]. Notably, incubation of VSV with a monoacylglycerol lipid form of C18:1 reduced its ability to inhibit infectivity, demonstrating that free FAs are more active against enveloped viruses than lipids [24]. Unlike these earlier studies, we found that HCMV replication through viral genome synthesis was unaffected or minimally impacted. Further work is needed to determine if EA directly targets HCMV, potentially disrupting its membrane, or if virus replication is reduced through an indirect mechanism, such as stimulation of intrinsic antiviral responses. Nonetheless, our work demonstrates that ELOVL5 reduces EA inhibition of HCMV. We propose that HCMV infection elevates ELOLV5 levels and activity to increase the pool of VLC-PUFAs (Figure 10B). Increasing the pool of VLC-PUFAs will support lipid synthesis, including the sequestration of EA into lipids. Overall, our study reveals that virus reprogramming of metabolism may increase the production of an antiviral metabolite or FA that can be further metabolized to evade its antiviral effects.

**Figure 10.**
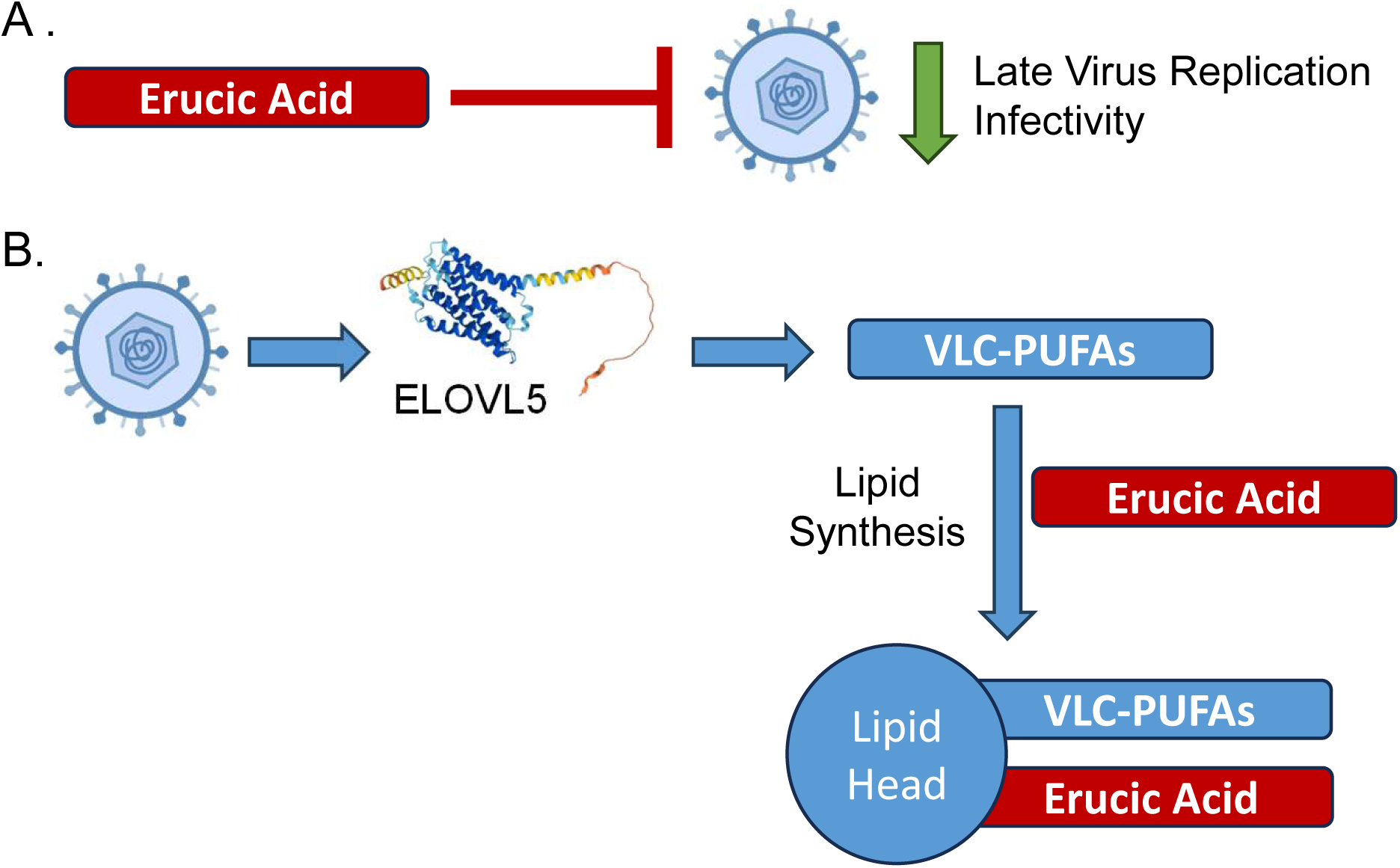
Proposed model for ELOVL5 activity in metabolizing EA to lipids in HCMV infection. (A) EA suppresses HCMV by targeting the late replication stage and reducing virus progeny infectivity. (B) HCMV infection promotes ELOVL5 levels to increase the synthesis of VLC-PUFAs. EA inhibition of HCMV is suppressed by feeding EA and VLC-PUFAs into lipid synthesis to generate EA containing lipids. AlphaFold rendering of human ELOVL5 (AF-Q9NYP7-F1).

## MATERIALS AND METHODS

### Cells and Viruses

Human foreskin fibroblast (HFF) cells were cultured in Dulbecco’s modified Eagle’s medium (DMEM) containing 10% fetal bovine serum (FBS), 10 mM HEPES, and penicillin-streptomycin. Prior to infection, cells were maintained at full confluence for 3 days in serum-containing growth medium. Cells were switched to serum-free medium (DMEM, HEPES, and penicillin-streptomycin) the day before infection. HCMV AD169 or TB40/E-GFP strains were used throughout the study [23, 53–55]. Cells were infected at a multiplicity of infection (MOI) 1 or 3 infectious units per cell, as indicated. Uninfected cells were mock-infected by treating the cells in the same manner as infected cells, except that the inoculum lacked virus particles. At 48 hpi, the growth medium was replaced with fresh media to maintain nutrient levels. All virus stocks were made by pelleting virus from the supernatant of infected cells through 20% sorbitol using ultracentrifugation. Viruses were resuspended in serum-free DMEM and stored at-80°C. Infectious virus titers were measured by 50% tissue culture infectious dose (TCID_50_). Particle-to-infectious unit ratio was determined by calculating the ratio between viral DNA and infectious virus yield in cell-free supernatants. Determination of virus yields from cell-free supernatants is described elsewhere [56]. Viral DNA in supernatants was quantified by qPCR as previously described [57].

For particle-to-IU ratio, 100 µL of culture supernatants from 120 and 144 hpi were incubated with 5 Units of DNase I (ThermoFisher) for 1 h at 37°C. DNA was isolated using Quick DNA mini-Prep Plus kit (Zymo). Primers to UL123 and an absolute standard curve using BAC-Fix-Actin were used to determine HCMV particle DNA levels.

### Generation of ELOVL5 Knockout Cells Using CRISPR/Cas9 Engineering

sgRNA sequence specific for human gene ELOVL was cloned into LentiCRISPR-v2 [58, 59], which co-expresses a mammalian codon-optimized Cas9 nuclease and a sgRNA [45]. A sgRNA that does not target any human or HCMV sequence is used for a non-targeting (NT) control. The gRNA sequences used in CRISPR knockout experiments are: ELOVL5-KO, AACAGCCATTCTCTTGCCGG; NT, CGCTTCCGCGGCCCGTTCAA.

Lentivirus expressing Cas9 and sgRNA were produced in 293T cells and transduced into life-extended primary human fibroblasts (HFF-hTERT), followed by drug selection using 2 µg/ml puromycin. Single-cell clones were obtained by diluting transduced cells into a 96-well plate, along with 200 non-transduced HFF cells, as previously described [45]. CRISPR-treated cells were selected a second time using 2 µg/ml puromycin once the cells reached 85-90% confluence. ELOVL5 knockout was confirmed by western blot and sequencing using the Guide-IT Indel kit (Takara Bio) and primers targeting the corresponding genomic region of the ELOVL5 gene. Each clone was sequenced at least 20 times to ensure biallelic mutations in the ELOVL5 gene. In parallel, the ELOVL5 gene was sequenced in the NT cells to confirm that the HFF-hTERT cells used contained a WT ELOVL5 gene.

### Analysis of nucleic acid

Viral and cellular DNA and RNA were determined by quantitative PCR (qPCR) using Power SYBR Green PCR master mix (ThermoFisher) and QuantStudio 3 Flex real-time PCR instrument. DNA isolation was completed using Quick DNA mini-Prep Plus kit (Zymo). An absolute standard curve made by diluting a BAC containing the HCMV strain FIX and actin sequence and primers to HCMV UL123 and cellular actin was used to determine relative viral to cellular genome levels. RNA isolation was performed using Quick-RNA Mini Prep Kit (Zymo), and cDNA was made using Transcriptor First Strand cDNA Synthesis Kit (Roche). Relative HCMV UL123, UL122, UL44, UL32, UL99, and UL82 to H6PD gene levels were determined by delta delta CT method. The sequences of primers are listed in the following table:

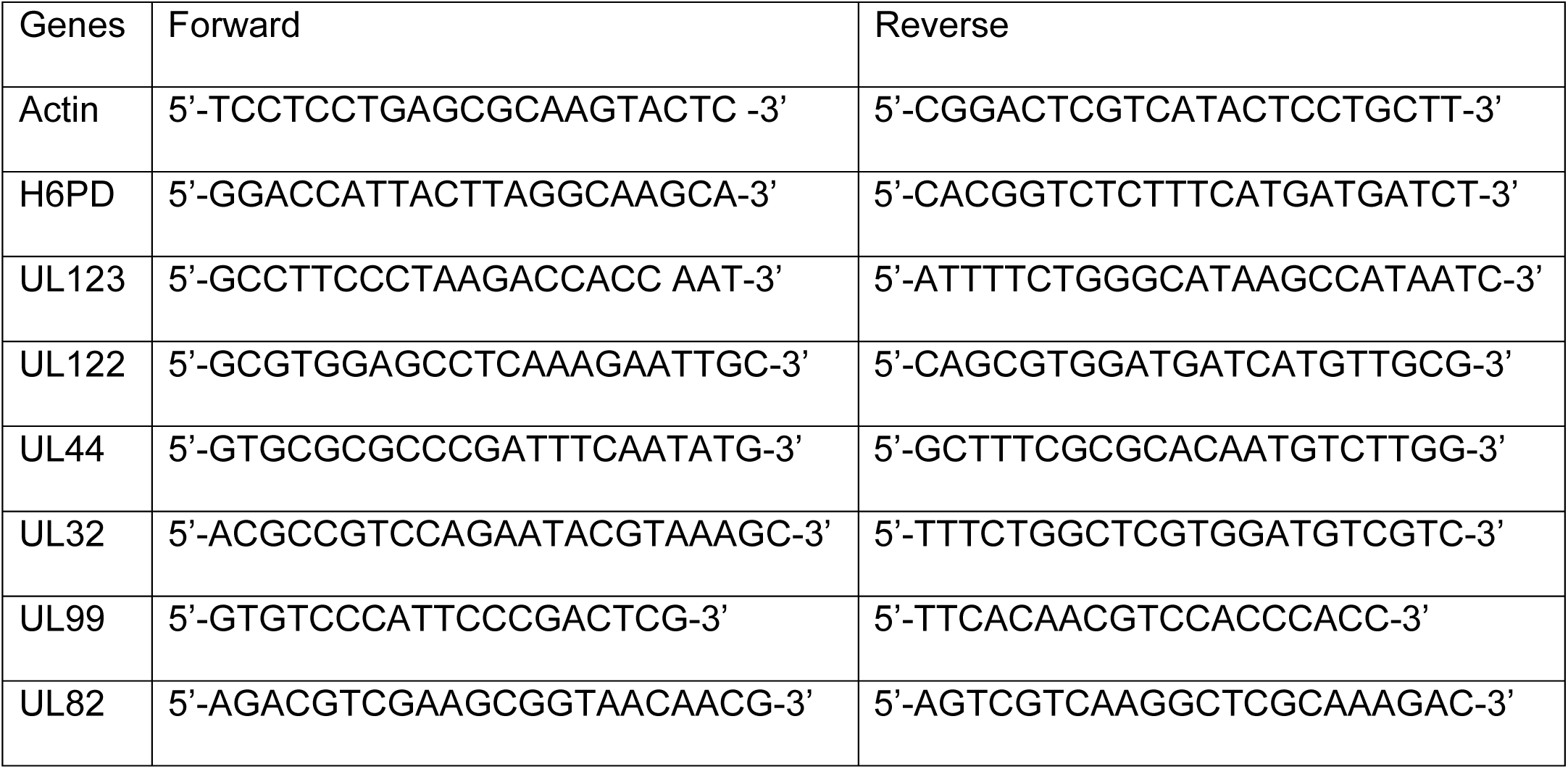

### Erucic acid (EA) treatment

HFF or ELOVL5-KO cells were grown to 100% confluency and serum-starved for 24 h before infection. Cells were infected at MOI 1. EA was conjugated to FA-free bovine serum albumin (BSA, Sigma A8806) before addition to cells. BSA stock solutions were prepared in 10 mM NaCl. BSA was diluted in DMEM for a final concentration of 2.5%. EA stocks were prepared in 100% ethanol. Thus, ethanol was used as the vehicle control throughout the paper. The final concentration of ethanol is 0.5%. EA or ethanol was added to the BSA-DMEM mixture and incubated for 10 mins. Growth media with EA or ethanol was added to the cells at 1 hpi, after the initial inoculum was removed and the cells were washed with warm PBS. EA was replenished at 48 hpi. The toxicity of EA on HFFs was determined via LDH Cytotoxicity Assay (Invitrogen).

### Lipid Extractions

For cellular lipids, HFF cells were grown to 100% confluency and serum-starved for 24 hours before infection. Cells were infected at MOI 3. Lipids were extracted at 72 hpi. Cells were washed with PBS prior to analysis. Pre-chilled 50% methanol was added to the cells, and lipids were extracted twice using chloroform. The chloroform was removed by drying the samples under a stream of nitrogen gas. Lipids were resuspended in a 1:1:1 solution of methanol, chloroform, and isopropanol. For each sample, 3 wells on a 6-well plate were used. Lipids were extracted in parallel from two wells and independently analyzed (e.g., technical duplicates). The cells in the third well were counted. Samples were normalized according to the total number of live cells at the time of lipid extraction. Each sample was resuspended in 100 μl of a 1:1:1 solution per 200,000 cells. In parallel with the biological samples, the extraction procedure was performed on wells that lacked cells. These “no-cell” controls were used to determine contaminants that were removed from the analysis.

### Lipidomics

The abundance of lipids was measured using liquid chromatography–high-resolution tandem mass spectrometry (LC-MS/MS) as previously described [23]. Briefly, cells were washed with PBS and lysed in cold 50% methanol. Lipids were extracted twice using chloroform and then dried under a stream of nitrogen gas. Lipids then were resuspended in 100 µl of a 1:1:1 solution of methanol-chloroform-isopropanol per 100,000 cells. For each sample, a total of three wells were used for analysis. Two wells were used for lipid extraction (i.e., duplicate samples analyzed in parallel to determine technical variation), and one well was used to determine the total number of cells.

Samples were normalized according to the number of live cells at the time of lipid extraction. A fourth well with no cells was used as a control to determine if any contaminants were introduced during the lipid extraction and LC-MS/MS steps. Any mass spectral feature from this contaminants list that appeared in the samples was removed from the analysis. During data collection, the samples were stored at 4°C in an autosampler.

Splash Lipidmix lipidomic mass spectrometry standards (Avanti Polar Lipids) were used to determine extraction efficiencies and quantify lipids. Following resuspension, lipids were stored at 4 °C in an autosampler. Lipids were separated by reverse-phase chromatography using a Kinetex 2.6 µm C18 column (Phenomenex 00F-4462-AN). LC was performed at 60 °C using a Vanquish UHPLC system (Thermo Scientific) with two solvents: solvent A (40:60 water-methanol, plus 10 mM ammonium formate and 0.1% formic acid) and solvent B (10:90 methanol-isopropanol, plus 10 mM ammonium formate and 0.1% formic acid). UHPLC was performed at a flow rate of 0.25 mL/min, starting at 25% solvent B and ending at 100% solvent B, as described [23].

After each sample, the column was washed and equilibrated. The total runtime was 30 min per sample. Blank samples were run before, after, and interspersed with the samples. Lipids were measured using a Q-Exactive Plus mass spectrometer operating in a data-dependent full MS/dd-MS2 TopN mode. MS1 data were collected at a resolution of either 70,000 or 140,000. Additional settings included an automatic gain control (AGC) target of 1e6, transient times of 250 ms for a 70,000 resolution, and 520 ms for a 140,000 resolution. MS1 Spectra were collected over a mass range of 200 to 1,600 m/z. MS2 spectra were collected using a transient time of 120 ms and a resolution setting of 35,000 with an AGC target of 1e5. Each sample was analyzed using negative and positive ion modes. The mass analyzer was calibrated weekly.

Lipids were ionized using a heated electrospray ionization (HESI) source and nitrogen gas as described [23]. Lipids were identified and quantified using MAVEN [60], EI-MAVEN (Elucidata), and Xcalibur (Thermo Scientific), as previously described [23].

### Protein Analysis

Proteins were examined by western blot using SDS-PAGE performed with tris-glycine-SDS running buffer. Proteins were separated using Mini-Protean TGX anyKD or 4 to 20% gels (Bio-Rad) and transferred to an Odyssey nitrocellulose membrane (Li-Cor). Membranes were blocked using 5% milk in tris-buffered saline with 0.05% Tween 20 (TBS-T) and incubated with primary antibodies in the presence of 1% milk TBS-T solution for anti-ELOVL5. For HCMV primary antibodies, membranes were blocked using 5% BSA in TBS-T and incubated with primary antibodies in the presence of 3% BSA in TBS-T. The following antibodies were used: mouse monoclonal anti-pUL123 (IE1, 1:50 dilution), mouse monoclonal anti-pUL44 (Virusys; ICP36; 1:2,500 dilution), mouse monoclonal anti-pUL26 (1:100 dilution), mouse monoclonal anti-pUL99 (pp28; 1:100 dilution), mouse monoclonal anti-pUL83 (pp71; 1:100 dilution), rabbit polyclonal anti-ELOVL7 (Sigma-Aldrich; #SAB3500390; 1:1,000 dilution), rabbit polyclonal anti-ELOVL5 (Sigma-Aldrich; #SAB4502642; 1:500 dilution), rabbit polyclonal anti-β-actin (Proteintech; #20536-1-AP; 1:2,000 dilution), and mouse monoclonal anti-α-tubulin (Sigma-Aldrich; #T6199; 1:2,000 dilution). Blots with mouse monoclonal anti-HCMV, anti-actin, and anti-tubulin antibodies were incubated for 1h at room temperature. All others were incubated overnight at 4°C. Quantification of western blots was performed using a Li-Cor Odyssey CLx imaging system.

### NF-κB measurement

HFF cells were infected with TB40/E at MOI 1 followed by EA treatment at 0, 5, 25, and 50 µM at 1 hpi. Cells were collected at 72 hpi. Separation of nuclear and cytosolic fractions was performed using a cellular nuclear extraction kit, according to the manufacturer’s instructions (Cayman; 10009277). The purity of cytosolic and nuclear fractions was confirmed by western blot staining for tubulin and lamin B, respectively. The nuclear fraction was used for testing NF-κB activation using the NF-κB (p65) transcription factor assay kit according to the manufacturer’s instructions (Cayman; #10007889). Briefly, pure nuclear fraction was added into a 96-well plate that was pre-coated with a consensus dsDNA sequence containing the NF-κB response element. A primary antibody against NF-κB (p65) and a secondary antibody conjugated to HRP were used for the detection of NF-κB (p65). The absorbance was measured at 450 nm. A positive control (a clarified cell lysate for NF-κB activation provided in the kit) and negative controls were used in this assay.

## Acknowledgements

We thank Nate Schmidt and Dr. Debbie Mustacich for providing technical assistance that contributed to the development of this project. Further, we are grateful to Dr. Felicia Goodrum, Dr. Sam Campos, Dr. James Alwine, and the other members of the Purdy lab for providing helpful feedback on this project. We would also like to thank Dr. Stanley Lemon for his pioneering work on HAV that has demonstrated the importance of studying lipids with VLCFA tails.

This project was supported by the following National Institute of Allergy and Infectious Disease (NIAID), National Institutes of Health (NIH) awards: R01AI162671 (J.G.P.), R01AI155539 (J.G.P.), and F32AI178919 (R.L.M.). Additional support was provided from the NIH National Institute of Aging (NIA) T32AG058503 award (I.K.). The content is solely the responsibility of the authors and does not necessarily represent the official views of the NIH. Additional funding was provided by the Personalized Defenses Against Disease strategic initiative from the University of Arizona in support X.Y. and the BIO5 Institute Postdoctoral Fellowship program in support of R.L.M. Finally, additional support for this work was provided by a New Investigator Award (ADHS18-198868) to J.G.P. from the Arizona Biomedical Research Commission (ABRC), made available through the Arizona Department of Health Services.

---

**Figure S1.**
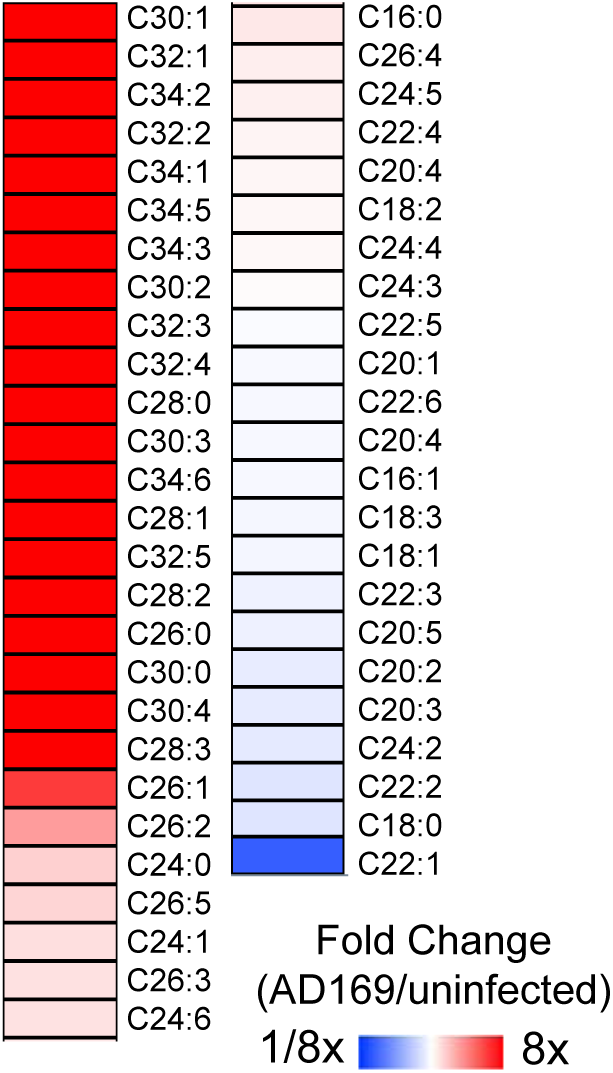
FA profile of AD169 infected cells. FA analysis of uninfected and HCMV AD169 infected HFF cells. Relative fold change of infected to uninfected. MOI 3, 72 hpi. N=3

**Figure S2.**
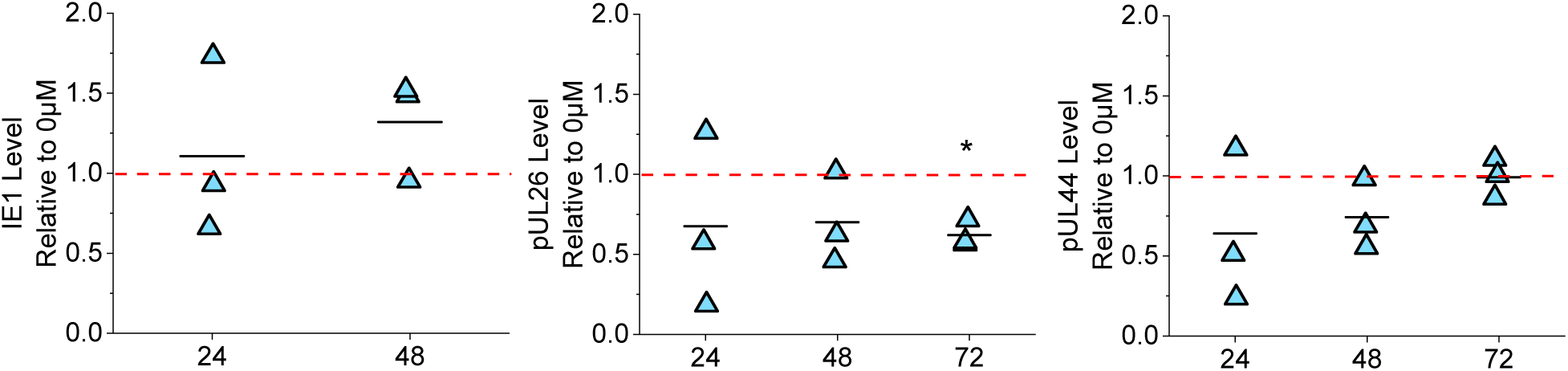
Levels of immediate-early and early viral genes in EA treatment. Quantification of immediate early (IE1) and early (pUL26, pUL44) viral protein from western blot shown in Fig 2A. Protein levels were normalized to 0 μM EA treated at each time. Red dash line represents 1-fold. MOI 1. N=3. One sample t-test, *p<0.05

**Figure S3.**
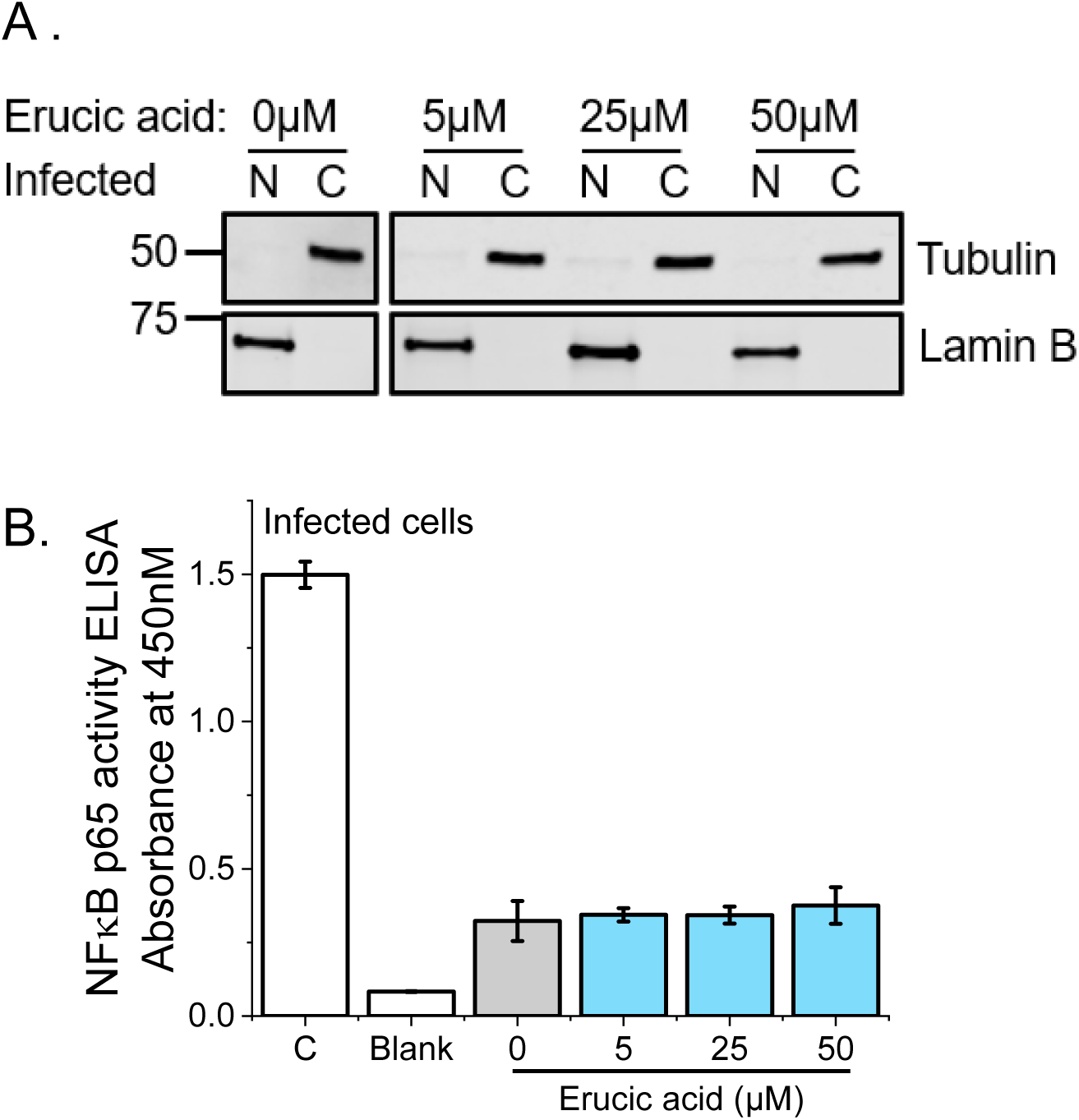
EA does not activate NFκB. (A) Western blot of protein in EA treated HCMV-infected cells. The nuclear and cytoplasmic fractions were separated. Blot for tubulin and lamin B as indicators of cytosolic and nuclear fraction, respectively. Tubulin and lamin B were blot on the same gel. (B) ELISA of NFκB p65 activity in EA treated infected cells in nuclear fraction compared to 0 μM. A positive control (C) that contains a clarified cell lysate for NFκB activation and a negative control (Blank) that only has assay binding buffer were included for comparison. N=3

**Figure S4.**
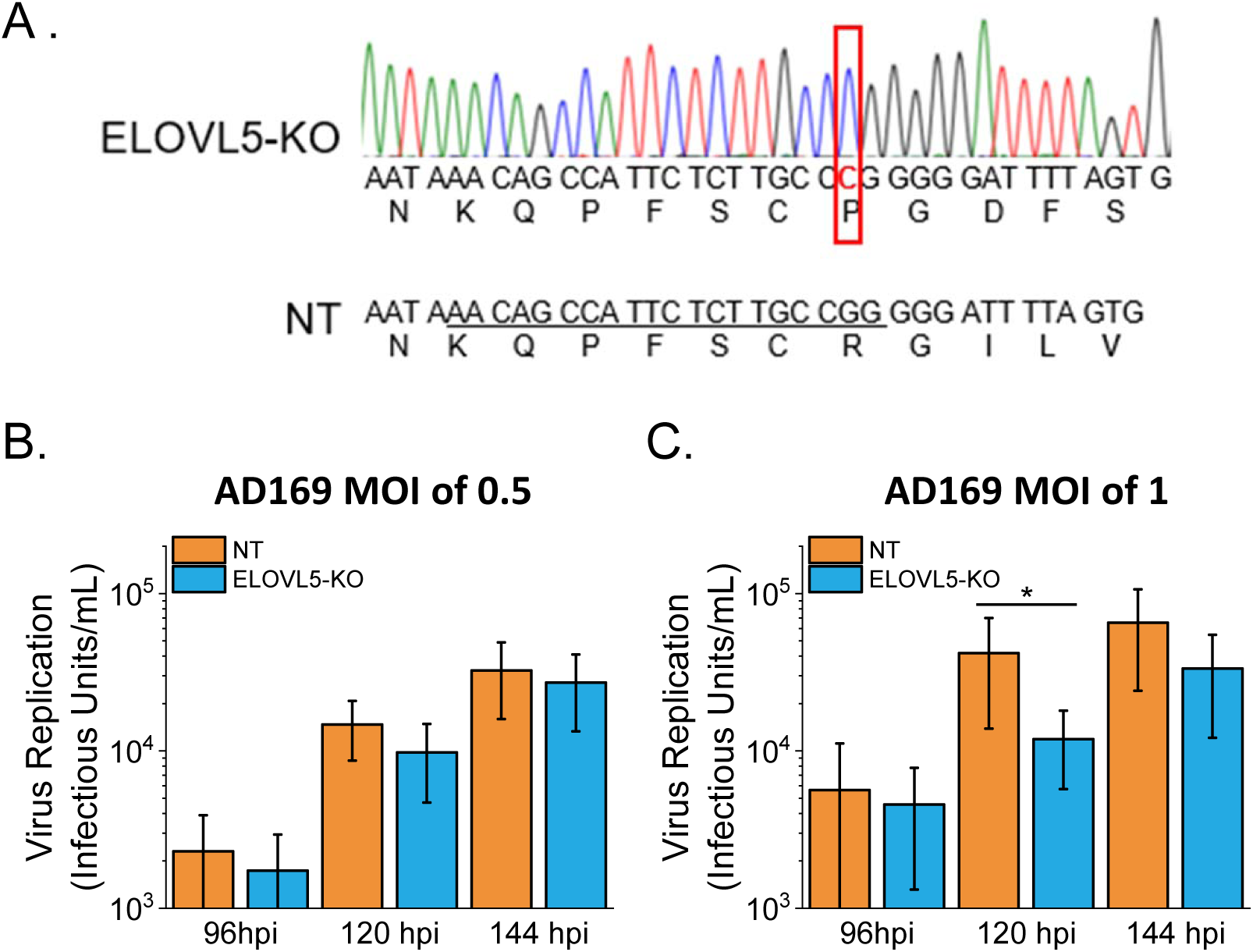
ELOVL5 is not required for HCMV replication. (A) Chromatogram of ELOVL5 CRISPR knockout (ELOVL5-KO) sequence and non-targeting (NT) control sequence (WT ELOVL5). A cytosine insertion occurs leading to a frame shift and loss of ELOVL5 protein sequence. Insertion is marked in red box. (B) Virus replication in AD169-infected NT and ELOVL5-KO HFF-hTERT cells at 96, 120, and 144 hpi measured by TCID50. N=3

**Figure S5.**
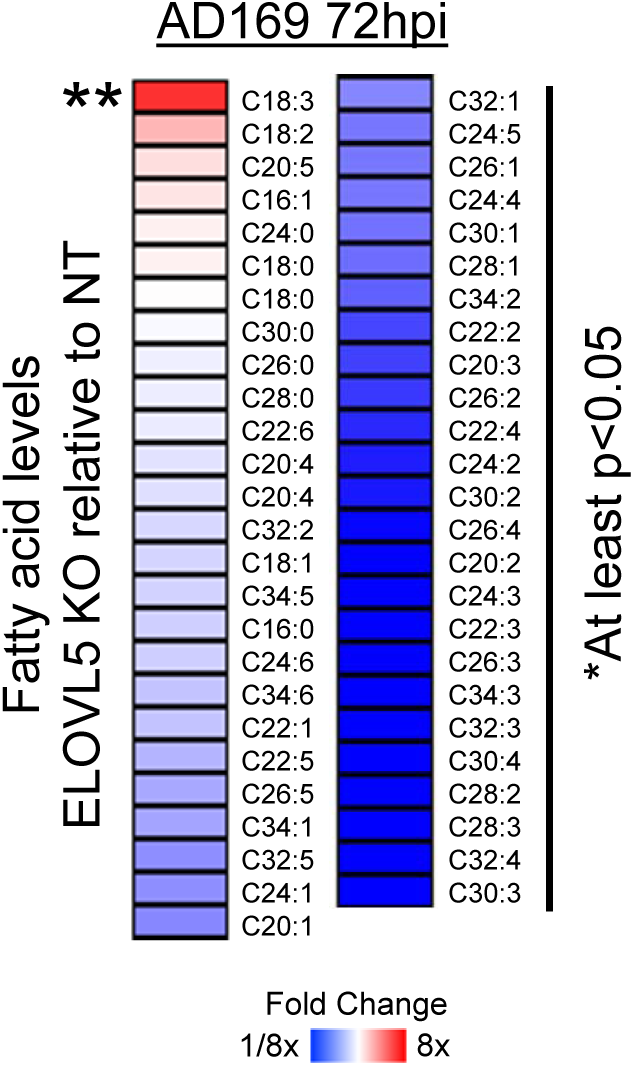
Depletion of ELOVL5 reduces VLC-PUFAs and EA. FA levels in AD169-infected ELOVL5-KO cells relative to infected NT cells. N=3. One sample t-test was used. *p>0.05, **p<0.01, ***p<0.001

**Figure S6.**
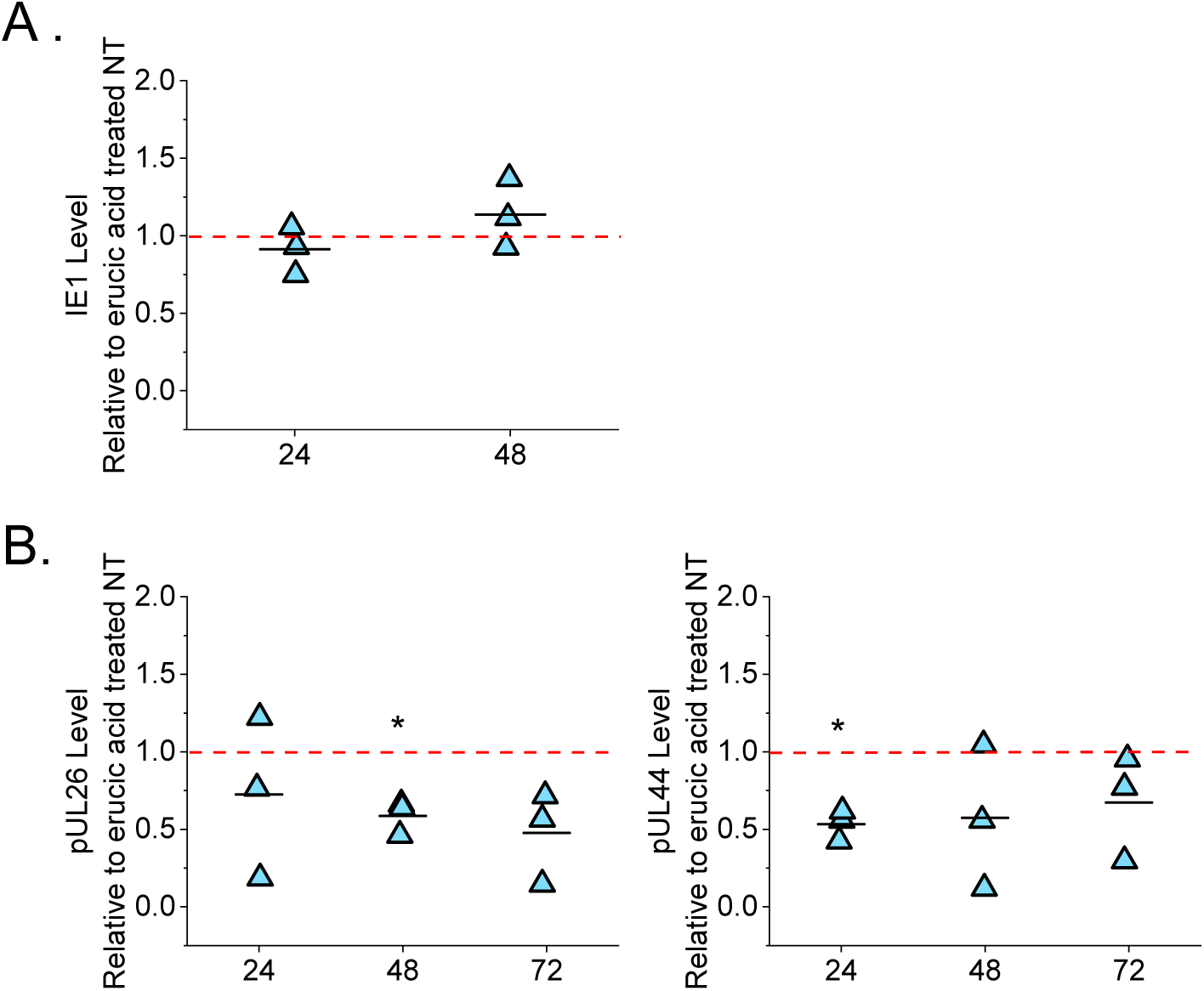
Immediate early and early viral protein levels in EA treated ELOVL-5 KO cells. (A) Quantification of IE1 in 50 μM EA treated ELOVL5-KO cells relative to NT cells from Fig 7A. (B) Quantification of early (pUL26 and pUL44) viral protein levels in 50 μM erucic acid treated ELOVL5-KO cells relative to NT cells from Fig 7A. N=3. One sample t-test *p<0.053

**Table S1.** Fatty acid composition in lipids that were downregulated by EA treatment.

| Lipid | Observed Tails | Fold Difference relative to uninfected 0 μM EA | | | |
| --- | --- | --- | --- | --- | --- |
|  |  | Uninfected |  | Infected |  |
| | | 0 μM | 50 μM | 0 μM | 50 μM |
| PE(36:6) | C14:0, C16:0, C16:1, C18:0, C20:4, C20:5, C22:6 | 1 | 0.1 | 1.5 | 0.1 |
| PS(42:1) | **C22:1**, C16:1, C18:1, C24:0 | 1 | 0.2 | 1.1 | 0.1 |
| PG(32:0) | C16:0 | 1 | 0.1 | 1 | 0.1 |
| PS(44:4) | C18:1, C20:4, C22:4, C24:1 | 1 | 0.3 | 1 | 0.3 |
| PE(48:7) | **C22:1**, C20:4, C22:4, C22:5, C22:6, C24:1, C26:1 | 1 | 0.2 | 1.1 | 0.2 |
| PC(28:1) | C14:0, C14:1, C16:0, C16:1 | 1 | 0.3 | 4.1 | 0.2 |
| PE(36:5) | C14:0, C16:0, C16:1, C18:0, C18:2, C20:4, C20:5 | 1 | 0.3 | 1.1 | 0.2 |
| PE(32:2) | C16:0, C16:1, C16:2 | 1 | 0.4 | 1 | 0.3 |
| PE(34:3) | C16:0, C16:1, C16:2, C18:0, C18:1, C18:2, C18:3, C20:3 | 1 | 0.4 | 1.1 | 0.3 |
| PE(38:6) | C16:0, C16:1, C18:0, C18:1, C18:2, C20:4, C20:5, C22:6 | 1 | 0.4 | 0.9 | 0.4 |
| PC(28:0) | C14:0, C16:0 | 1 | 0.4 | 4.8 | 0.2 |
| PG(38:6) | C16:0, C16:1, C18:0, C18:1, C20:5, C22:5, C22:6 | 1 | 0.4 | 0.9 | 0.4 |
| PS(38:6) | C16:0, C22:6 | 1 | 0.4 | 0.9 | 0.3 |
| PC(48:5) | C20:4, C22:5, C28:1 | 1 | 0.4 | 5.6 | 0.7 |
| PC(30:2) | C14:0, C14:1, C16:0, C16:1 | 1 | 0.4 | 3.1 | 0.3 |
| PG(32:2) | C16:0, C16:1, C18:1 | 1 | 0.5 | 1.3 | 0.6 |
| PG(38:7) | **C22:1**, C16:1, C18:0 | 1 | 0.5 | 0.9 | 0.7 |
| PE(36:4) | C16:0, C16:1, C18:1, C18:2, C18:3, C20:3, C20:4 | 1 | 0.5 | 1.1 | 0.4 |
| PG(44:12) | C18:0, C22:6 | 1 | 0.5 | 0.8 | 0.6 |
| PE(32:0) | C14:0, C16:0, C18:0 | 1 | 0.5 | 1.9 | 0.3 |
| PE(32:1) | C14:0, C16:0, C16:1, C18:1 | 1 | 0.5 | 1.4 | 0.5 |
| PG(32:1) | C14:0, C16:0, C16:1, C18:1 | 1 | 0.5 | 1.3 | 0.4 |
| PC(30:1) | C14:0, C16:1 | 1 | 0.5 | 1.9 | 0.4 |
| PC(42:0) | C16:0, C18:1, C26:0 | 1 | 0.6 | 2.4 | 0.5 |
| PG(44:11) | C18:0, C22:5, C22:6 | 1 | 0.6 | 0.6 | 0.6 |
| PC(36:5) | C16:0, C16:1, C16:2, C18:0, C18:1, C18:2, C18:3, C18:4, C20:4, C20:5, C22:5 | 1 | 0.6 | 0.8 | 0.4 |
| PE(34:4) | C14:0, C16:0, C16:1, C16:2, C18:1, C18:2, C18:3, C20:4 | 1 | 0.6 | 1.7 | 0.6 |
| PC(30:0) | C14:0, C16:0 | 1 | 0.6 | 1.2 | 0.4 |
| PC(34:4) | C14:0, C16:0, C16:1, C18:0, C18:1, C18:2, C18:3, C20:4 | 1 | 0.6 | 1.2 | 0.4 |
| PC(48:6) | C22:0 | 1 | 0.6 | 4.4 | 0.9 |
| PE(38:0) | C16:0, C16:1, C18:0, C20:0, C22:0 | 1 | 0.6 | 0.9 | 0.5 |
| PC(36:4) | C16:0, C16:1, C16:2, C18:0, C18:1, C18:2, C18:3, C20:3, C20:4 | 1 | 0.6 | 0.9 | 0.4 |
| PC(34:3) | C14:0, C16:0, C16:1, C16:2, C18:1, C18:2, C18:3, C20:3 | 1 | 0.6 | 1 | 0.5 |
| PE(36:0) | C16:0, C18:0 | 1 | 0.6 | 0.9 | 0.5 |

C22:1 FA tails are shown in bold.

**Table S2.**
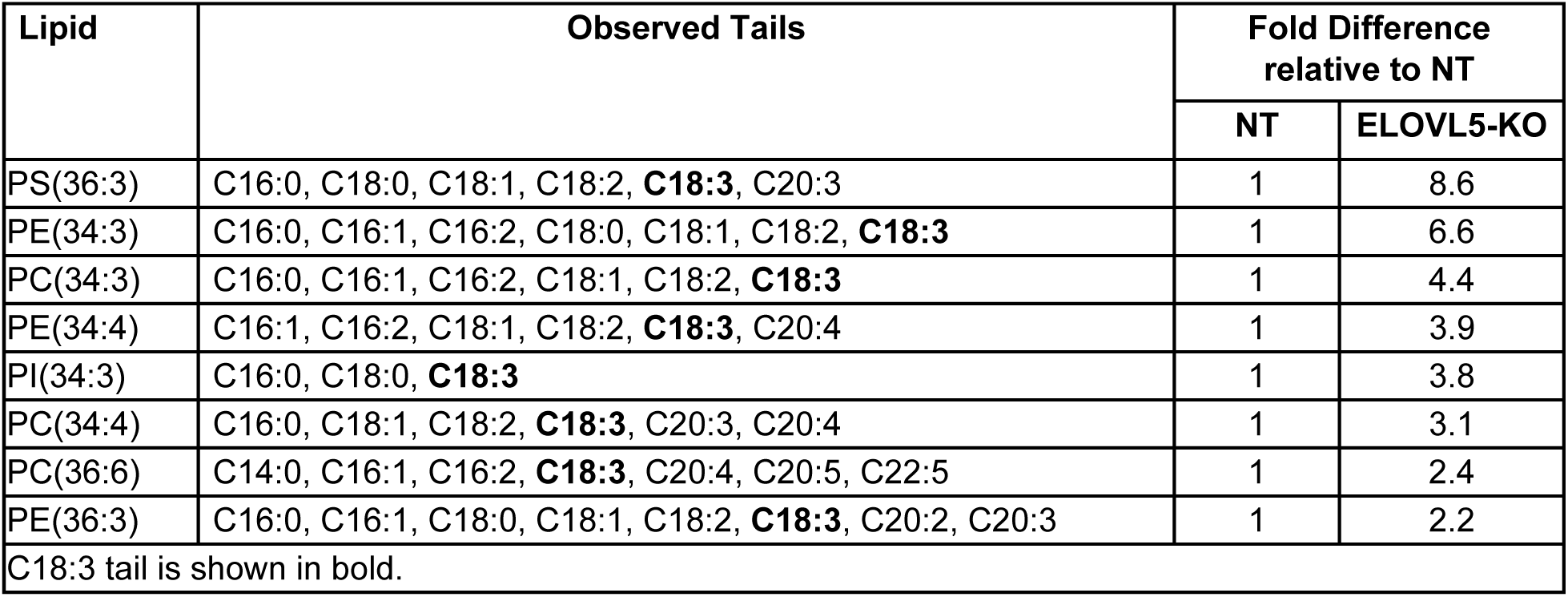
FA composition of lipids upregulated in HCMV-infected ELOVL5-KO cells.

**Table S3.**
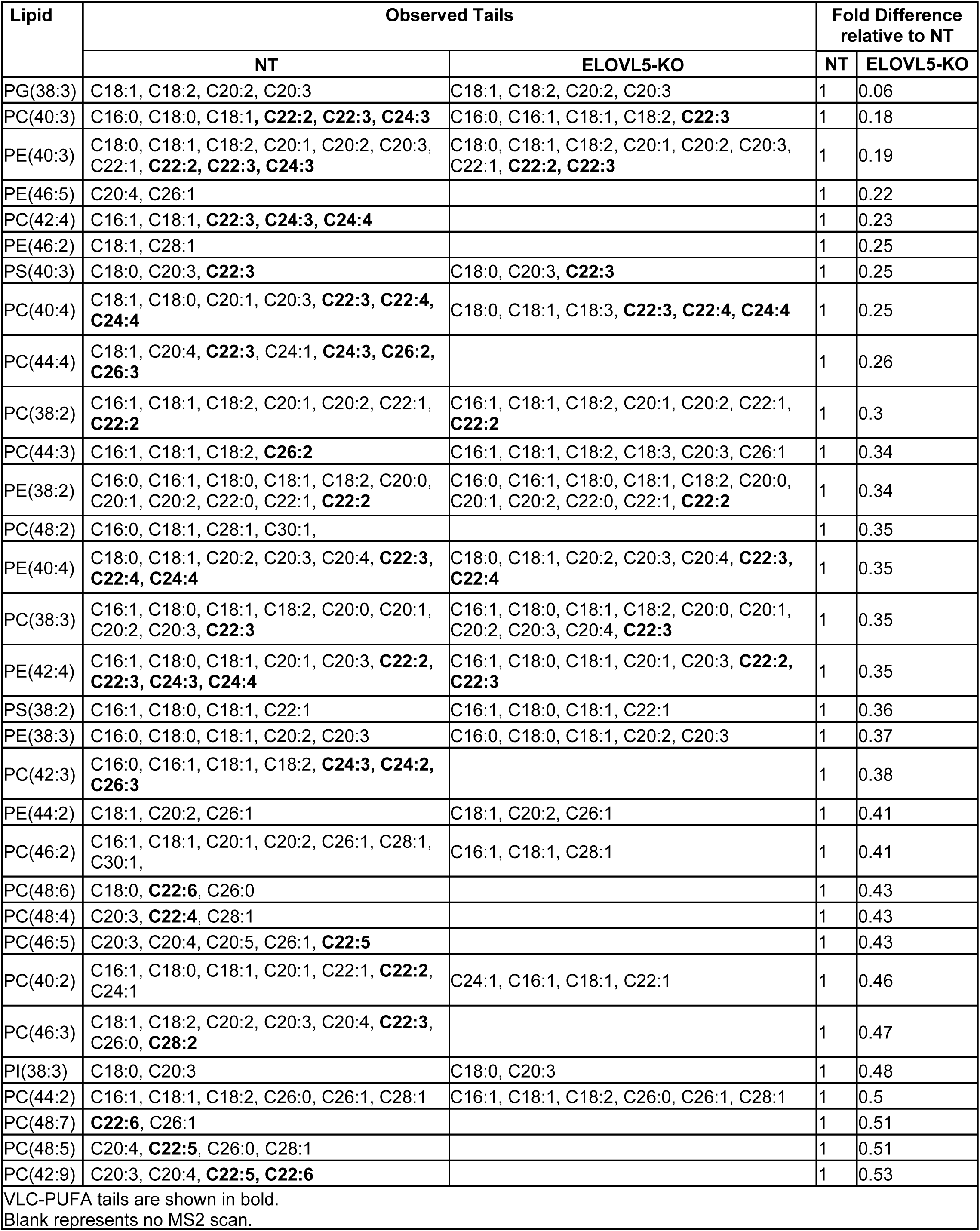
FA composition of lipids downregulated in HCMV-infected ELOVL5-KO cells.

## REFERENCES

1. Cannon MJ, Schmid DS, Hyde TB. Review of cytomegalovirus seroprevalence and demographic characteristics associated with infection. Rev Med Virol. 2010;20(4):202–13. doi: 10.1002/rmv.655. PubMed PMID: 20564615.

2. Mocarski ES, Shenk T, Pass RF, editors. Cytomegaloviruses. Philadelphia, PA: Lippincott, Williams and Wilkins; 2007.

3. Britt W. Manifestations of human cytomegalovirus infection: proposed mechanisms of acute and chronic disease. Current topics in microbiology and immunology. 2008;325:417–70. PubMed PMID: 18637519.

4. Kenneson A, Cannon MJ. Review and meta-analysis of the epidemiology of congenital cytomegalovirus (CMV) infection. Rev Med Virol. 2007;17(4):253–76. Epub 2007/06/21. doi: 10.1002/rmv.535. PubMed PMID: 17579921.

5. Lanzieri TM, Dollard SC, Bialek SR, Grosse SD. Systematic review of the birth prevalence of congenital cytomegalovirus infection in developing countries. Int J Infect Dis. 2014;22:44–8. Epub 20140312. doi: 10.1016/j.ijid.2013.12.010. PubMed PMID: 24631522; PubMed Central PMCID: PMCPMC4829484.

6. Lischka P, Zimmermann H. Antiviral strategies to combat cytomegalovirus infections in transplant recipients. Current opinion in pharmacology. 2008;8(5):541–8. Epub 2008/07/30. doi: 10.1016/j.coph.2008.07.002. PubMed PMID: 18662804.

7. Biron KK. Antiviral drugs for cytomegalovirus diseases. Antiviral Res. 2006;71(2-3):154–63. Epub 2006/06/13. doi: 10.1016/j.antiviral.2006.05.002. PubMed PMID: 16765457.

8. Xi Y, Lindenmayer L, Kline I, von Einem J, Purdy JG. Human Cytomegalovirus Uses a Host Stress Response To Balance the Elongation of Saturated/Monounsaturated and Polyunsaturated Very-Long-Chain Fatty Acids. mBio. 2021;12(3). doi: 10.1128/mBio.00167-21. PubMed PMID: 33947752.

9. Xi Y, Harwood S, Wise L, Purdy JG. Human Cytomegalovirus pUL37×1 is Important to Remodeling of Host Lipid Metabolism. J Virol. 2019:JVI.00843-19. doi: 10.1128/jvi.00843-19.

10. Purdy JG, Shenk T, Rabinowitz JD. Fatty Acid Elongase 7 Catalyzes Lipidome Remodeling Essential for Human Cytomegalovirus Replication. Cell reports. 2015;10(8):1375–85. Epub 2015/03/04. doi: 10.1016/j.celrep.2015.02.003. PubMed PMID: 25732827; PubMed Central PMCID: PMCPMC4354725.

11. Koyuncu E, Purdy JG, Rabinowitz JD, Shenk T. Saturated very long chain fatty acids are required for the production of infectious human cytomegalovirus progeny. PLoS pathogens. 2013;9(5):e1003333. Epub 2013/05/23. doi: 10.1371/journal.ppat.1003333. PubMed PMID: 23696731; PubMed Central PMCID: PMC3656100.

12. Ohno Y, Suto S, Yamanaka M, Mizutani Y, Mitsutake S, Igarashi Y, et al. ELOVL1 production of C24 acyl-CoAs is linked to C24 sphingolipid synthesis. Proc Natl Acad Sci U S A. 2010;107(43):18439–44. doi: 10.1073/pnas.1005572107. PubMed PMID: 20937905; PubMed Central PMCID: PMCPMC2973002.

13. Moon YA, Hammer RE, Horton JD. Deletion of ELOVL5 leads to fatty liver through activation of SREBP-1c in mice. Journal of lipid research. 2009;50(3):412–23. Epub 2008/10/08. doi: 10.1194/jlr.M800383-JLR200. PubMed PMID: 18838740; PubMed Central PMCID: PMC2638104.

14. Tamura K, Makino A, Hullin-Matsuda F, Kobayashi T, Furihata M, Chung S, et al. Novel lipogenic enzyme ELOVL7 is involved in prostate cancer growth through saturated long-chain fatty acid metabolism. Cancer research. 2009;69(20):8133–40. Epub 2009/10/15. doi: 0008-5472.CAN-09-0775 [pii]10.1158/0008-5472.CAN-09-0775. PubMed PMID: 19826053.

15. Naganuma T, Sato Y, Sassa T, Ohno Y, Kihara A. Biochemical characterization of the very long-chain fatty acid elongase ELOVL7. FEBS letters. 2011;585(20):3337–41. Epub 2011/10/01. doi: 10.1016/j.febslet.2011.09.024. PubMed PMID: 21959040.

16. Yu Y, Pierciey FJ, Jr., Maguire TG, Alwine JC. PKR-like endoplasmic reticulum kinase is necessary for lipogenic activation during HCMV infection. PLoS pathogens. 2013;9(4):e1003266. doi: 10.1371/journal.ppat.1003266. PubMed PMID: 23592989; PubMed Central PMCID: PMCPMC3617203.

17. McArdle J, Moorman NJ, Munger J. HCMV targets the metabolic stress response through activation of AMPK whose activity is important for viral replication. PLoS pathogens. 2012;8(1):e1002502. Epub 2012/02/01. doi: 10.1371/journal.ppat.1002502. PubMed PMID: 22291597; PubMed Central PMCID: PMC3266935.

18. Spencer CM, Schafer XL, Moorman NJ, Munger J. Human cytomegalovirus induces the activity and expression of acetyl-coenzyme A carboxylase, a fatty acid biosynthetic enzyme whose inhibition attenuates viral replication. J Virol. 2011;85(12):5814–24. Epub 2011/04/08. doi: 10.1128/JVI.02630-10. PubMed PMID: 21471234; PubMed Central PMCID: PMC3126312.

19. Raymonda MH, Rodríguez-Sánchez I, Schafer XL, Smorodintsev-Schiller L, Harris IS, Munger J. Cytomegalovirus-induced inactivation of TSC2 disrupts the coupling of fatty acid biosynthesis to glucose availability resulting in a vulnerability to glucose starvation. mBio. 2023:e0303123. Epub 20231220. doi: 10.1128/mbio.03031-23. PubMed PMID: 38117060.

20. Yu Y, Maguire TG, Alwine JC. Human cytomegalovirus infection induces adipocyte-like lipogenesis through activation of sterol regulatory element binding protein 1. J Virol. 2012;86(6):2942–9. Epub 2012/01/20. doi: 10.1128/JVI.06467-11. PubMed PMID: 22258239; PubMed Central PMCID: PMC3302344.

21. Betsinger CN, Jankowski CSR, Hofstadter WA, Federspiel JD, Otter CJ, Jean Beltran PM, et al. The human cytomegalovirus protein pUL13 targets mitochondrial cristae architecture to increase cellular respiration during infection. Proc Natl Acad Sci U S A. 2021;118(32). doi: 10.1073/pnas.2101675118. PubMed PMID: 34344827; PubMed Central PMCID: PMCPMC8364163.

22. Rodriguez-Sanchez I, Schafer XL, Monaghan M, Munger J. The Human Cytomegalovirus UL38 protein drives mTOR-independent metabolic flux reprogramming by inhibiting TSC2. PLoS pathogens. 2019;15(1):e1007569. doi: 10.1371/journal.ppat.1007569. PubMed PMID: 30677091; PubMed Central PMCID: PMCPMC6363234.

23. Xi Y, Harwood S, Wise L, Purdy JG. Human Cytomegalovirus pUL37×1 is Important to Remodeling of Host Lipid Metabolism. J Virol. 2019;93(21):1–19. Epub Aug 7. doi: 10.1128/JVI.00843-19. PubMed PMID: 31391267; PubMed Central PMCID: PMCPMC6803270.

24. Thormar H, Isaacs CE, Brown HR, Barshatzky MR, Pessolano T. Inactivation of enveloped viruses and killing of cells by fatty acids and monoglycerides. Antimicrobial agents and chemotherapy. 1987;31(1):27–31. doi: 10.1128/AAC.31.1.27. PubMed PMID: 3032090; PubMed Central PMCID: PMCPMC174645.

25. Stock CC, Francis T. The Inactivation of the Virus of Epidemic Influenza by Soaps. J Exp Med. 1940;71(5):661–81. doi: 10.1084/jem.71.5.661. PubMed PMID: 19870990; PubMed Central PMCID: PMCPMC2135101.

26. Kohn A, Gitelman J, Inbar M. Interaction of polyunsaturated fatty acids with animal cells and enveloped viruses. Antimicrobial agents and chemotherapy. 1980;18(6):962–8. Epub 1980/12/01. PubMed PMID: 7235682; PubMed Central PMCID: PMC352998.

27. Kohn A, Gitelman J, Inbar M. Unsaturated free fatty acids inactivate animal enveloped viruses. Archives of virology. 1980;66(4):301–7. Epub 1980/01/01. PubMed PMID: 7447706.

28. Liang X, Huang Y, Pan X, Hao Y, Chen X, Jiang H, et al. Erucic acid from Isatis indigotica Fort. suppresses influenza A virus replication and inflammation in vitro and in vivo through modulation of NF-kappaB and p38 MAPK pathway. J Pharm Anal. 2020;10(2):130–46. Epub 20191004. doi: 10.1016/j.jpha.2019.09.005. PubMed PMID: 32373385; PubMed Central PMCID: PMCPMC7192973.

29. Cappa M, Bizzarri C, Petroni A, Carta G, Cordeddu L, Valeriani M, et al. A mixture of oleic, erucic and conjugated linoleic acids modulates cerebrospinal fluid inflammatory markers and improve somatosensorial evoked potential in X-linked adrenoleukodystrophy female carriers. Journal of Inherited Metabolic Disease. 2012;35(5):899–907. doi: 10.1007/s10545-011-9432-3.

30. Munger J, Bajad SU, Coller HA, Shenk T, Rabinowitz JD. Dynamics of the cellular metabolome during human cytomegalovirus infection. PLoS pathogens. 2006;2(12):e132. Epub 2006/12/19. doi: 10.1371/journal.ppat.0020132. PubMed PMID: 17173481; PubMed Central PMCID: PMC1698944.

31. Munger J, Bennett BD, Parikh A, Feng XJ, McArdle J, Rabitz HA, et al. Systems-level metabolic flux profiling identifies fatty acid synthesis as a target for antiviral therapy. Nat Biotechnol. 2008;26(10):1179–86. Epub 2008/09/30. doi: nbt.1500 [pii]10.1038/nbt.1500. PubMed PMID: 18820684; PubMed Central PMCID: PMC2825756.

32. Vastag L, Koyuncu E, Grady SL, Shenk TE, Rabinowitz JD. Divergent effects of human cytomegalovirus and herpes simplex virus-1 on cellular metabolism. PLoS pathogens. 2011;7(7):e1002124. Epub 2011/07/23. doi: 10.1371/journal.ppat.1002124. PubMed PMID: 21779165; PubMed Central PMCID: PMC3136460.

33. Hwang J, Purdy JG, Wu K, Rabinowitz JD, Shenk T. Estrogen-related receptor alpha is required for efficient human cytomegalovirus replication. Proc Natl Acad Sci U S A. 2014;111(52):E5706-15. Epub 2014/12/17. doi: 10.1073/pnas.1422361112. PubMed PMID: 25512541; PubMed Central PMCID: PMC4284536.

34. Sun X, Song L, Feng S, Li L, Yu H, Wang Q, et al. Fatty Acid Metabolism is Associated With Disease Severity After H7N9 Infection. EBioMedicine. 2018;33:218–29. Epub 20180623. doi: 10.1016/j.ebiom.2018.06.019. PubMed PMID: 29941340; PubMed Central PMCID: PMCPMC6085509.

35. Liang X, Huang Y, Pan X, Hao Y, Chen X, Jiang H, et al. Erucic acid from Isatis indigotica Fort. suppresses influenza A virus replication and inflammation in vitro and in vivo through modulation of NF-κB and p38 MAPK pathway. Journal of Pharmaceutical Analysis. 2020;10(2):130–46. doi: 10.1016/j.jpha.2019.09.005.

36. Guillou H, Zadravec D, Martin PG, Jacobsson A. The key roles of elongases and desaturases in mammalian fatty acid metabolism: Insights from transgenic mice. Progress in lipid research. 2010;49(2):186–99. Epub 2009/12/19. doi: 10.1016/j.plipres.2009.12.002. PubMed PMID: 20018209.

37. Mokry RL, Purdy JG. Glucose-independent human cytomegalovirus replication is supported by metabolites that feed upper glycolytic branches. Proc Natl Acad Sci U S A. 2024;121(48):e2412966121. Epub 20241119. doi: 10.1073/pnas.2412966121. PubMed PMID: 39560652; PubMed Central PMCID: PMCPMC11621781.

38. Purdy JG, Luftig MA. Reprogramming of cellular metabolic pathways by human oncogenic viruses. Current opinion in virology. 2019;39:60–9. doi: 10.1016/j.coviro.2019.11.002. PubMed PMID: 31766001.

39. Rodriguez-Sanchez I, Munger J. Meal for Two: Human Cytomegalovirus-Induced Activation of Cellular Metabolism. Viruses. 2019;11(3). doi: 10.3390/v11030273. PubMed PMID: 30893762; PubMed Central PMCID: PMCPMC6466105.

40. Lange PT, Lagunoff M, Tarakanova VL. Chewing the Fat: The Conserved Ability of DNA Viruses to Hijack Cellular Lipid Metabolism. Viruses. 2019;11(2). doi: 10.3390/v11020119. PubMed PMID: 30699959; PubMed Central PMCID: PMCPMC6409581.

41. Thaker SK, Ch’ng J, Christofk HR. Viral hijacking of cellular metabolism. BMC Biol. 2019;17(1):59. doi: 10.1186/s12915-019-0678-9. PubMed PMID: 31319842; PubMed Central PMCID: PMCPMC6637495.

42. Pant A, Dsouza L, Yang Z. Alteration in Cellular Signaling and Metabolic Reprogramming during Viral Infection. mBio. 2021;12(5):e0063521. doi: 10.1128/mBio.00635-21. PubMed PMID: 34517756; PubMed Central PMCID: PMCPMC8546648.

43. Sumbria D, Berber E, Mathayan M, Rouse BT. Virus Infections and Host Metabolism-Can We Manage the Interactions? Front Immunol. 2020;11:594963. doi: 10.3389/fimmu.2020.594963. PubMed PMID: 33613518; PubMed Central PMCID: PMCPMC7887310.

44. McGettrick AF, O’Neill LA. Two for the price of one: itaconate and its derivatives as an anti-infective and anti-inflammatory immunometabolite. Curr Opin Immunol. 2023;80:102268. Epub 20221126. doi: 10.1016/j.coi.2022.102268. PubMed PMID: 36446152.

45. Vysochan A, Sengupta A, Weljie AM, Alwine JC, Yu Y. ACSS2-mediated acetyl-CoA synthesis from acetate is necessary for human cytomegalovirus infection. Proc Natl Acad Sci U S A. 2017;114(8):E1528-E35. Epub 20170206. doi: 10.1073/pnas.1614268114. PubMed PMID: 28167750; PubMed Central PMCID: PMCPMC5338361.

46. Xi Y, Lindenmayer L, Kline I, von Einem J, Purdy JG. Human Cytomegalovirus uses a Host Stress Response to Balance the Elongation of Saturated/Monounsaturated and Polyunsaturated Very Long Chain Fatty Acids. bioRxiv. 2021:2021.01.16.426974. doi: 10.1101/2021.01.16.426974.

47. Yu Y, Maguire TG, Alwine JC. Human cytomegalovirus activates glucose transporter 4 expression to increase glucose uptake during infection. J Virol. 2011;85(4):1573–80. Epub 2010/12/15. doi: 10.1128/JVI.01967-10. PubMed PMID: 21147915; PubMed Central PMCID: PMC3028904.

48. Hofmann S, Krajewski M, Scherer C, Scholz V, Mordhorst V, Truschow P, et al. Complex lipid metabolic remodeling is required for efficient hepatitis C virus replication. Biochim Biophys Acta Mol Cell Biol Lipids. 2018;1863(9):1041–56. doi: 10.1016/j.bbalip.2018.06.002. PubMed PMID: 29885363.

49. Roe B, Kensicki E, Mohney R, Hall WW. Metabolomic profile of hepatitis C virus-infected hepatocytes. PLoS One. 2011;6(8):e23641. Epub 2011/08/20. doi: 10.1371/journal.pone.0023641. PubMed PMID: 21853158; PubMed Central PMCID: PMCPMC3154941 Inc. This does not alter the authors’ adherence to all of the PLoS ONE policies on sharing data and materials.

50. Douglas DN, Pu CH, Lewis JT, Bhat R, Anwar-Mohamed A, Logan M, et al. Oxidative Stress Attenuates Lipid Synthesis and Increases Mitochondrial Fatty Acid Oxidation in Hepatoma Cells Infected with Hepatitis C Virus*. Journal of Biological Chemistry. 2016;291(4):1974–90. doi: 10.1074/jbc.M115.674861.

51. Hofmann S, Krajewski M, Scherer C, Scholz V, Mordhorst V, Truschow P, et al. Complex lipid metabolic remodeling is required for efficient hepatitis C virus replication. Biochimica et Biophysica Acta (BBA) - Molecular and Cell Biology of Lipids. 2018;1863(9):1041–56. doi: 10.1016/j.bbalip.2018.06.002.

52. Shiota T, Li Z, Chen GY, McKnight KL, Shirasaki T, Yonish B, et al. Hepatoviruses promote very-long-chain fatty acid and sphingolipid synthesis for viral RNA replication and quasi-enveloped virus release. Sci Adv. 2023;9(42):eadj4198. Epub 20231020. doi: 10.1126/sciadv.adj4198. PubMed PMID: 37862421; PubMed Central PMCID: PMCPMC10588952.

53. Sinzger C, Hahn G, Digel M, Katona R, Sampaio KL, Messerle M, et al. Cloning and sequencing of a highly productive, endotheliotropic virus strain derived from human cytomegalovirus TB40/E. The Journal of general virology. 2008;89(Pt 2):359–68. Epub 2008/01/17. doi: 10.1099/vir.0.83286-0. PubMed PMID: 18198366.

54. Umashankar M, Petrucelli A, Cicchini L, Caposio P, Kreklywich CN, Rak M, et al. A novel human cytomegalovirus locus modulates cell type-specific outcomes of infection. PLoS pathogens. 2011;7(12):e1002444. Epub 2012/01/14. doi: 10.1371/journal.ppat.1002444. PubMed PMID: 22241980; PubMed Central PMCID: PMC3248471.

55. Yu D, Smith GA, Enquist LW, Shenk T. Construction of a self-excisable bacterial artificial chromosome containing the human cytomegalovirus genome and mutagenesis of the diploid TRL/IRL13 gene. J Virol. 2002;76(5):2316–28. Epub 2002/02/12. PubMed PMID: 11836410; PubMed Central PMCID: PMC153828.

56. Read C, Schauflinger M, Nikolaenko D, Walther P, von Einem J. Regulation of Human Cytomegalovirus Secondary Envelopment by a C-Terminal Tetralysine Motif in pUL71. J Virol. 2019;93(13):1–20. doi: 10.1128/JVI.02244-18. PubMed PMID: 30996102; PubMed Central PMCID: PMCPMC6580969.

57. Konig P, Svrlanska A, Read C, Feichtinger S, Stamminger T. The autophagy-initiating protein kinase ULK1 phosphorylates human cytomegalovirus tegument protein pp28 and regulates efficient virus release. J Virol. 2020. doi: 10.1128/JVI.02346-20. PubMed PMID: 33328309.

58. Sanjana NE, Shalem O, Zhang F. Improved vectors and genome-wide libraries for CRISPR screening. Nature methods. 2014;11(8):783–4. doi: 10.1038/nmeth.3047. PubMed PMID: 25075903; PubMed Central PMCID: PMCPMC4486245.

59. Shalem O, Sanjana NE, Hartenian E, Shi X, Scott DA, Mikkelson T, et al. Genome-scale CRISPR-Cas9 knockout screening in human cells. Science. 2014;343(6166):84-7. Epub 2013/12/18. doi: 10.1126/science.1247005. PubMed PMID: 24336571; PubMed Central PMCID: PMCPMC4089965.

60. Melamud E, Vastag L, Rabinowitz JD. Metabolomic analysis and visualization engine for LC-MS data. Analytical chemistry. 2010;82(23):9818–26. Epub 2010/11/06. doi: 10.1021/ac1021166. PubMed PMID: 21049934.

