## Supplemental Figures for "Human Cytomegalovirus Infection Reduces an Endogenous Antiviral Fatty Acid by Promoting Host Metabolism"

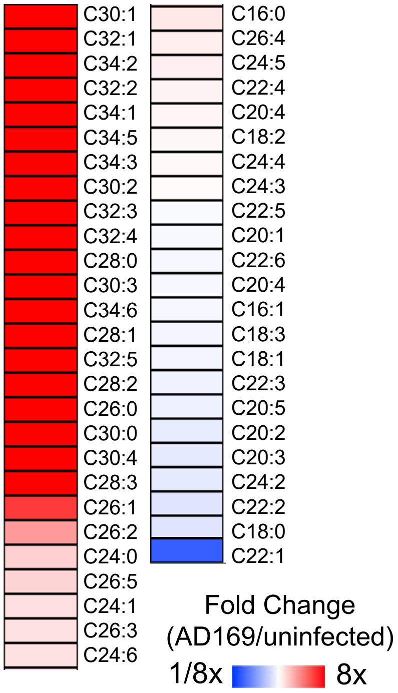


**Figure S1. FA profile of AD169 infected cells.** FA analysis of uninfected and HCMV AD169 infected HFF cells. Relative fold change of infected to uninfected. MOI 3, 72 hpi. N=3

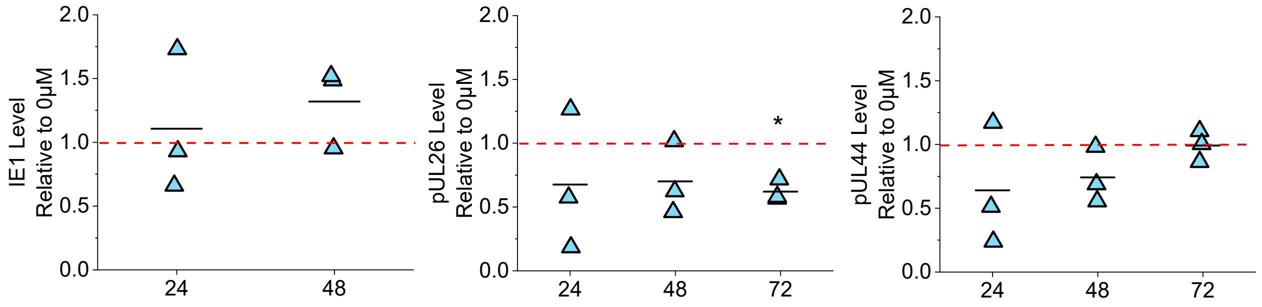


**Figure S2. Levels of immediate-early and early viral genes in EA treatment.** Quantification of immediate early (IE1) and early (pUL26, pUL44) viral protein from western blot shown in Fig 2A. Protein levels were normalized to 0 μM EA treated at each time. Red dash line represents 1-fold. MOI 1. N=3. One sample t-test, *p<0.05

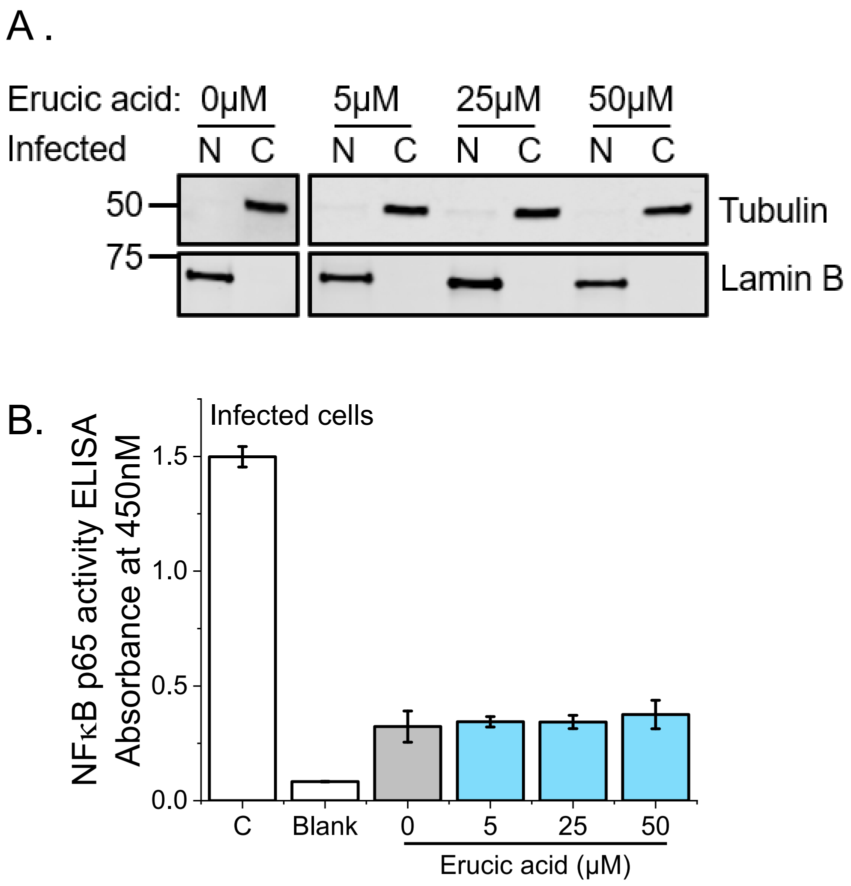


**Figure S3. EA does not activate NFκB.** (A) Western blot of protein in EA treated HCMV-infected cells. The nuclear and cytoplasmic fractions were separated. Blot for tubulin and lamin B as indicators of cytosolic and nuclear fraction, respectively. Tubulin and lamin B were blot on the same gel. (B) ELISA of NFκB p65 activity in EA treated infected cells in nuclear fraction compared to 0 μM. A positive control (C) that contains a clarified cell lysate for NFκB activation and a negative control (Blank) that only has assay binding buffer were included for comparison. N=3

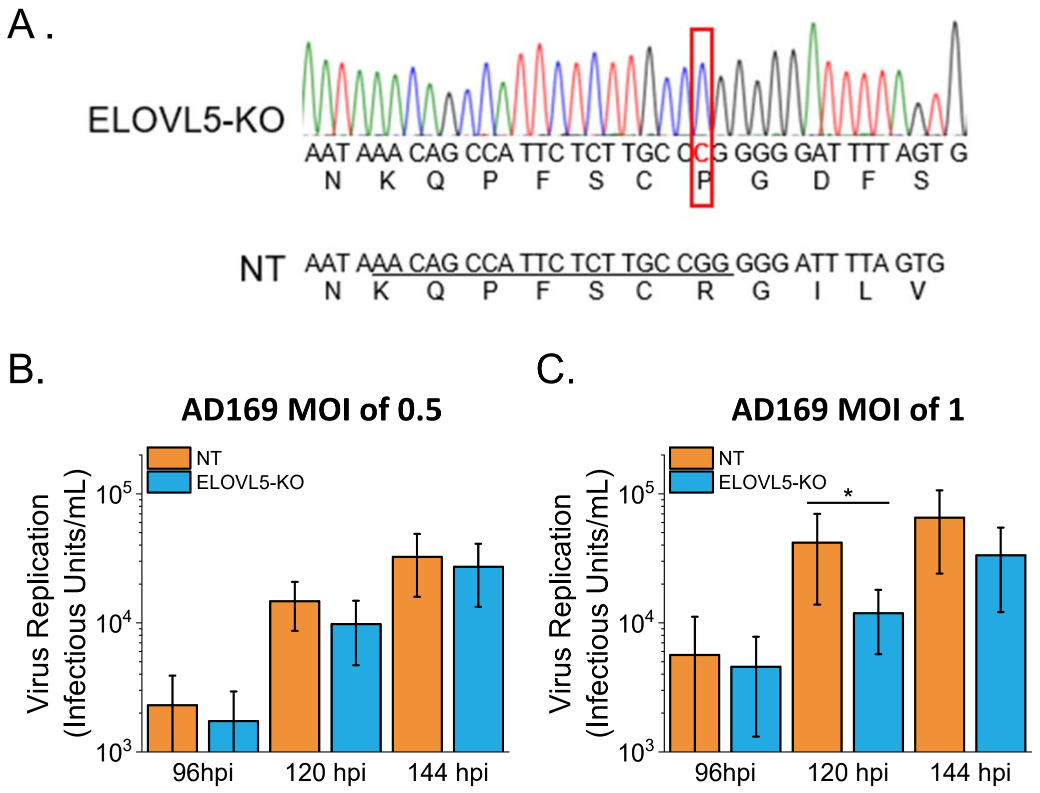


**Figure S4. ELOVL5 is not required for HCMV replication.** (A) Chromatogram of ELOVL5 CRISPR knockout (ELOVL5-KO) sequence and non-targeting (NT) control sequence (WT ELOVL5). A cytosine insertion occurs leading to a frame shift and loss of ELOVL5 protein sequence. Insertion is marked in red box. (B) Virus replication in AD169-infected NT and ELOVL5-KO HFF-hTERT cells at 96, 120, and 144 hpi measured by TCID50. N=3

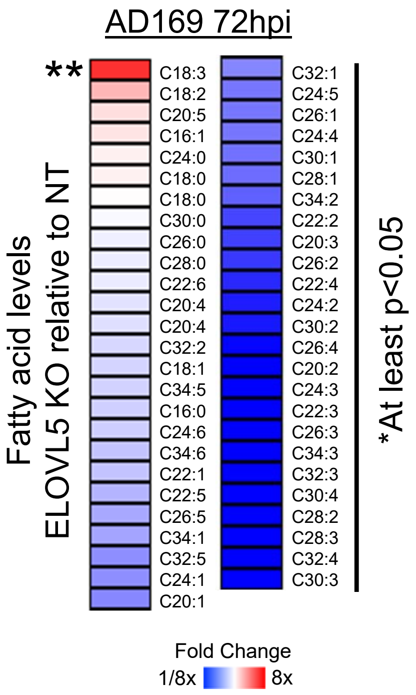


**Figure S5. Depletion of ELOVL5 reduces VLC-PUFAs and EA.** FA levels in AD169-infected ELOVL5-KO cells relative to infected NT cells. N=3. One sample t-test was used. *p>0.05, **p<0.01, ***p<0.001

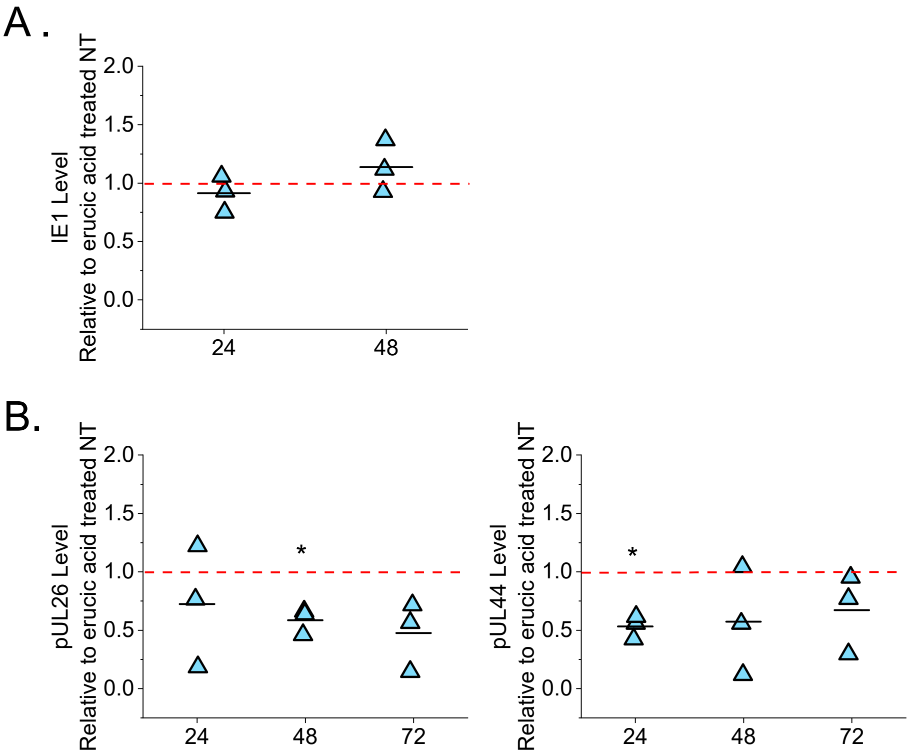


**Figure S6. Immediate early and early viral protein levels in EA treated ELOVL-5 KO cells.** (A) Quantification of IE1 in 50 μM EA treated ELOVL5-KO cells relative to NT cells from Fig 7A. (B) Quantification of early (pUL26 and pUL44) viral protein levels in 50 μM erucic acid treated ELOVL5-KO cells relative to NT cells from Fig 7A. N=3. One sample t-test $*p<0.053$
